# Efficient Inhibition of O-glycan biosynthesis using the hexosamine analog Ac_5_GalNTGc

**DOI:** 10.1101/2020.12.13.422539

**Authors:** Shuen-Shiuan Wang, Virginia del Solar, Xinheng Yu, Aristotelis Antonopoulos, Alan E. Friedman, Kavita Agarwal, Monika Garg, Syed Meheboob Ahmed, Ahana Addhya, Mehrab Nasirikenari, Joseph T. Lau, Anne Dell, Stuart M. Haslam, Srinivasa-Gopalan Sampathkumar, Sriram Neelamegham

**Author notes:** **Co-corresponding senior authors:** Sriram Neelamegham, 906 Furnas Hall, Buffalo, NY 14260, USA; or S.-Gopalan Sampathkumar, National Institute of Immunology, Aruna Asaf Ali Marg, New Delhi 110067 India. The authors contributed equally (Shuen-Shiuan Wang, Virginia del Solar).

## Abstract

There is a critical need to develop small molecule inhibitors of mucin-type O-linked glycosylation. The best known reagent currently is peracetylated benzyl-GalNAc, but it is only effective at millimolar concentrations. This manuscript demonstrates that Ac_5_GalNTGc, a peracetylated C-2 sulfhydryl substituted GalNAc, fulfills this unmet need. When added to cultured leukocytes, breast and prostate cells, Ac_5_GalNTGc increased cell surface VVA-binding by ~10-fold, indicating truncation of O-glycan biosynthesis. Cytometry, mass spectrometry and Western blot analysis of HL-60 promyelocytes demonstrate that 50-80μM Ac_5_GalNTGc prevented elaboration of 30-60% of the O-glycans beyond the Tn-antigen (GalNAcα1-Ser/Thr) stage. The effect of the compound on N-glycans and glycosphingolipids was small. Glycan inhibition induced by Ac_5_GalNTGc resulted in 50-80% reduction in leukocyte sialyl-Lewis-X expression, and L-/P-selectin mediated rolling under flow. Ac_5_GalNTGc was pharmacologically active in mouse. It reduced neutrophil infiltration to sites of inflammation by ~60%. Overall, Ac_5_GalNTGc may find diverse applications as a potent inhibitor of O-glycosylation.

## INTRODUCTION

All mammalian cells display mucin-type O-linked GalNAc-type glycans as part of their glycocalyx. They play a number of key biological roles during development, tumorigenesis, cancer metastasis, leukocyte adhesion and inflammatory response (Chugh et al., 2018; Oliveira-Ferrer et al., 2017; Schnaar, 2016; Tran and Ten Hagen, 2013). The biosynthesis of mucin-type O-glycans is initiated by the transfer of GalNAc (*N*-Acetylgalactosamine) from its nucleotide-sugar donor (UDP-GalNAc) to Ser/Thr residues on the peptide backbone by a family of polypeptide GalNAc-transferases (ppGalNAcT, (Bennett et al., 2012)). There are 20 ppGalNAcTs in humans, and these are largely conserved in the animal kingdom. Many of these enzymes act on proline-rich peptides to result in α-GalNAc-Ser/Thr (Tn-antigen) bearing glycopeptides. Other ‘follow-up enzymes’ (ppGalNAcT-T4, -T7, -T10, -T12, -T17) prefer to act on previously GalNAcylated glycopeptides to promote the formation of *O*-glycan clusters on mucinous proteins (Bennett et al., 2012). While the overlapping tissue expression patterns and substrate specificity of the ppGalNAcTs result in some functional redundancy, the second step of the *O*-glycan biosynthesis pathway is highly specific. Here, Galβ1→3 addition is mediated by a single, ubiquitously expressed enzyme called Core 1 β1,3Galactosyltransferase (C1GalT1) along with its chaperone protein Cosmc (Core 1 β1,3GalT-specific molecular chaperone) (Ju and Cummings, 2002). The Galβ1,3GalNAcα-Ser/Thr core is then elaborated by additional glycosyltransferases (Brockhausen and Stanley, 2015).

The development of specific, small molecule inhibitors of mucin-type O-glycans would be beneficial for mechanistic studies and also translational applications. This would fill a need in the field since many of the current glycosylation pathway inhibitors only target either glycosidase function (e.g. castanospermine, thiamet-G, oseltamivir), *N*-glycan biosynthesis (swainsonine, deoxymannojirimycin, tunicamycin), or glycosphingolipid (GSL) biosynthesis (*N*-butyldeoxynojirimycin; D-threo-1-phenyl-2-palmitoylamino-3-morpholino-l-propanol PPMP) (Gloster and Vocadlo, 2012; Hudak and Bertozzi, 2014). Attempts have been made to develop such *O*-glycosylation inhibitors with focus on GalNAc as these are uniquely part of *O*-linked glycans, although they also appear in dermatan sulfates, chondroitin sulfates and a subset of GSLs. Such compounds are often peracetylated to enhance cell permeability. In this regard, studies using various GalNAc analogs suggest that C-6 modification of GalNAc may render the compound inactive as it is not activated by GalNAc-1-kinase in the salvage pathway (Pouilly et al., 2012). However, C-2 *N*-acyl modified GalNAc are tolerated by cells and incorporated into cell surface glycoconjugates to varying degrees depending on their chemical composition (Dube et al., 2006; Hang et al., 2003; Pouilly et al., 2012). Such compounds are however reported to function bio-orthogonally, without inhibition effects. Peracetylated 4F-GalNAc is another C-4 analog that acts as a glycosylation inhibitor at 50-100μM concentrations (Marathe et al., 2010). Like peracetylated 4F-GlcNAc, however, this compound is not directly incorporated into glycoconjugates (Barthel et al., 2011), and both compounds reduce depress UDP-HexNAc levels (Del Solar et al., 2020). Finally, the most common *O*-glycan inhibitor used currently is the decoy substrate GalNAc-O-Bn (‘benzyl-α-GalNAc’) that acts as an effective surrogate substrate when applied at high concentrations (2-4mM) (Alfalah et al., 1999; Huet et al., 1998; Kuan et al., 1989; Tsuiji et al., 2003). At lower dose (25-100μM), peracetylated GalNAc-O-Bn only acts as a primer that reports on the cellular carbohydrate biosynthesis pathways with minimal inhibitory function (Kudelka et al., 2016; Stolfa et al., 2016).

Previously, we reported a peracetylated *N*-thioglycolyl modified GalNAc analog (‘Ac_5_GalNTGc’), which trimmed O-glycans globally, including on CD43 of Jurkat (Agarwal et al., 2013) and U937 cell lines (Dwivedi et al., 2018). In the current manuscript, we extended the characterization of this compound, with focus on its effect on leukocyte cell adhesion function, mechanisms of action and *in vivo* investigations of leukocyte recruitment. These studies contrast the function of Ac_5_GalNTGc with a panel of other peracetylated C-2 substituted GalNAc analogs, and also its peracetylated C-4 epimer, ‘Ac5GlcNTGc’ (**Figure 1**). These compounds are per-acetylated to enhance cell permeability. Once they enter cells from the culture medium, they are de-esterified in the cytosol and transported into the Golgi. Here, they either participate in biosynthetic processes or act as metabolic inhibitors. In such investigations, our data show that: i. Ac_5_GalNTGc effectively abolishes sialyl Lewis-X (sLe^X^) epitope expression on human leukocytes when applied at 50 μM, and inhibits leukocyte rolling on L- and P-selectin *ex vivo.* Such binding is largely dependent on leukocytic *O*-glycans (Kieffer et al., 2001; Lo et al., 2013; Vestweber and Blanks, 1999). Whereas the concentration in the extra-cellular milieu is in the ~50μM range, intra-cellular concentration is higher it ~1-1.5 mM (estimated in (Marathe et al., 2010)). ii. Addition of this compound to different cell types resulted in a dramatic upregulation of VVA-lectin binding, supporting the potential that Ac_5_GalNTGc acts by inhibiting core-1 glycan elaboration. iii. Although only a minor portion of the Ac_5_GalNTGc was incorporated into cellular glycoconjugates, upregulation of VVA-lectin binding correlated tightly with the extent of Ac_5_GalNTGc incorporation. Ac_5_GalNTGc did not alter cellular nucleotide-sugar levels or N-glycan biosynthesis. It had a quantitative effect on the relative abundance of GSLs. iv. Ac_5_GalNTGc was pharmacologically active in mice and it inhibited the extent of neutrophil recruitment to sites of inflammation in an acute peritonitis model. Overall, Ac_5_GalNTGc is a mucin-type O-glycosylation inhibitor. It is more potent compared to other molecules commonly used in literature, and thus could be broadly useful in diverse basic science and translational applications.

**Figure 1.**
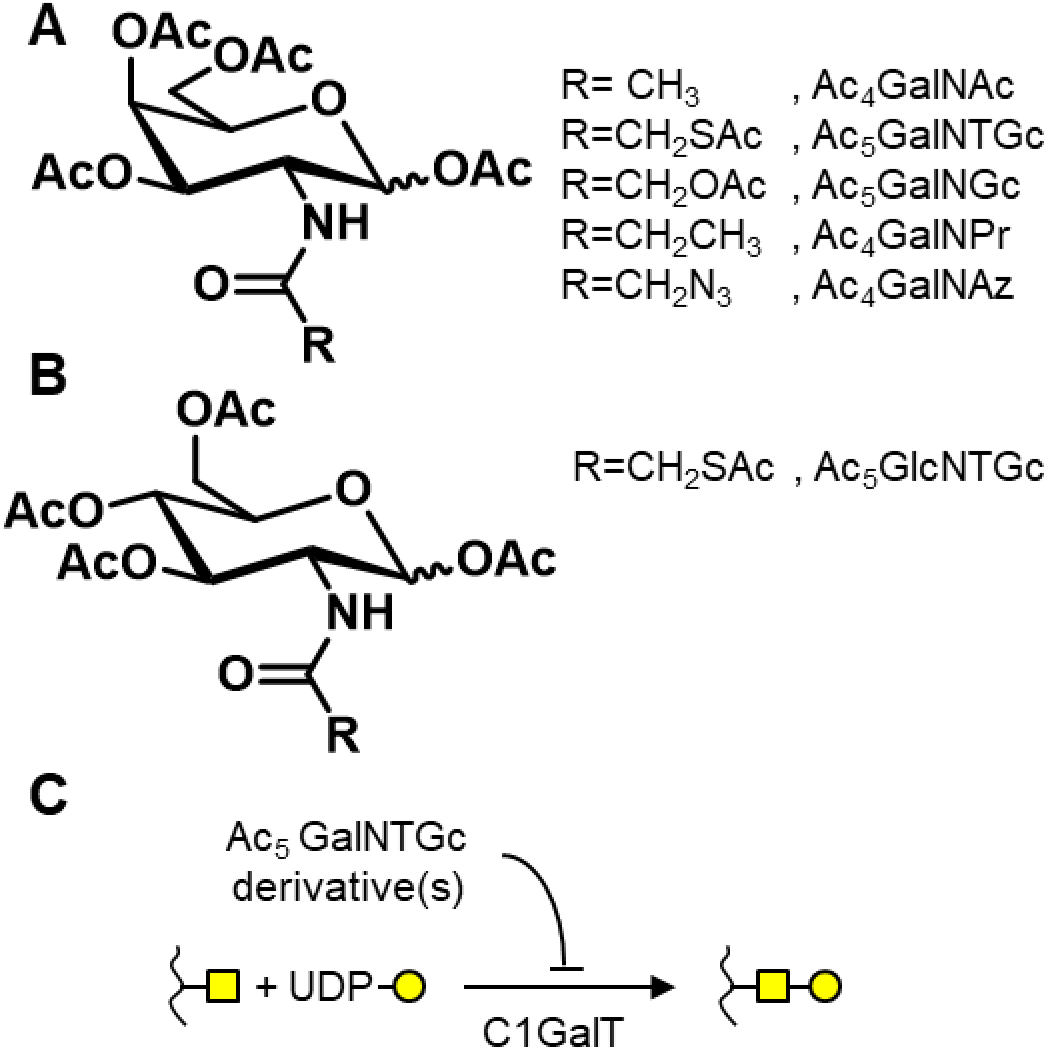
Peracetylated HexNAc analogs. **A.** Structures of per-acetylated GalNAc (Ac_4_GalNAc) and its analogs. **B.** Per-acetylated GlcNTGc (Ac_5_GlcNTGc), an isomer of peracetylated GalNTGc. **C.** Schematic showing the possible mechanism of Ac_5_GalNTGc action. Ac_5_GalNTGc or its derivatives inhibit GalNAc-type O-glycan biosynthesis.

## RESULTS

### Ac_5_GalNTGc decreased cell surface sialyl Lewis-X expression

The effect of the panel of peracetylated HexNAc analogs on cell surface carbohydrate expression was evaluated by culturing HL-60 promyelocytes with 50 μM of each of the analogs for 40h (**Fig. 2**). Among them, only Ac_5_GalNTGc significantly altered cell surface glycan structures. It doubled CD15/Le^X^ expression (Fig. 2A), reduced the expression of sLe^X^ as determined using mAbs HECA-452 (Fig. 2B) and CSLEX-1 by 50-80% (Fig. 2C), and increased the VIM-2/CD65s epitope (Fig. 2D). Other compounds containing different C-2 substituents, and also the C-4 epimer Ac_5_GlcNTGc had no effect on glycan expression. A reduction in HECA-452 (CLA/sLe^X^) expression, upon culture with Ac_5_GalNTGc, was also observed in western blots (Supplemental Fig. S1A). The increase in Le^X^ expression is similar to prior work where CRISPR-Cas9 based inhibition of O-glycan biosynthesis in HL-60s uncovered sterically hidden CD15 epitopes on leukocyte N-glycans and glycolipids (Stolfa et al., 2016). In dosage studies, Ac_5_GalNTGc modified glycan structures at concentrations as low as 10 μM, with maximum efficacy at > 50 μM (Fig. 2E-2H). None of the concentrations tested altered cell viability or growth rate over the first 40h, though treatment with 200 μM Ac_5_GalNTGc resulted in a longer lag-phase before resumption of growth following Ac_5_GalNTGc removal (Supplemental Fig. S1B-D). Overall, GalNAc with a thioglycolylamino-moiety at the C-2 position is a potent modifier of glycosylation. Based on these data, 80μM peracetylated Ac_5_GalNTGc was applied in all subsequent studies described below, unless stated otherwise.

**Figure 2.**
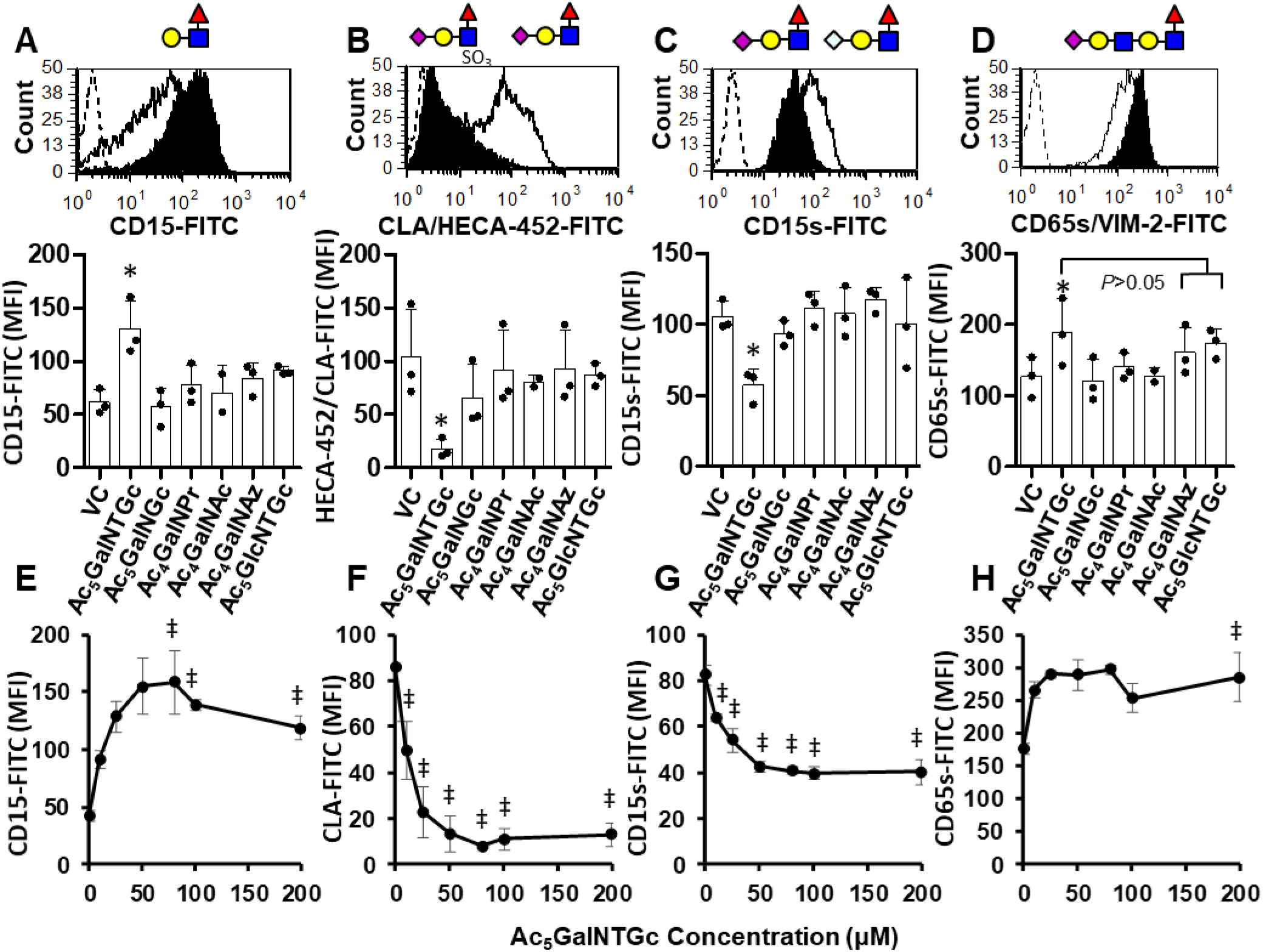
Effect of HexNAc analogs on cell-surface glycans. **A**-**D**. 0.5×10^6^ HL60 cells/mL were cultured with 50μM peracetylated GalNAc or GlcNAc analogs for 40h (VC=vehicle control). Cell-surface carbohydrate structures were measured using flow cytometry. Antigens measured include: **A.** CD15/Lewis-X (mAb HI98), **B.** Sialyl Lewis-X (sLe^X^) like antigen called CLA or Cutaneous Lymphocyte Antigen (mAb HECA-452), **C.** CD15s/sLe^X^ (mAb CSLEX-1) and **D.** the sialofucosylated epitope CD65s measured using mAb VIM-2. Flow cytometry histograms present data for isotype control (dashed-empty), VC (solid-empty) and Ac_5_GalNTGc (black-filled) treated samples. Bar plots present results for all analogs. \**P*<0.05 with respect to other treatments, except as indicated in panel D. **E**-**F**. Same studies as panel A-D, only Ac_5_GalNTGc concentration was titrated from 0-200μM. ‡ *P*<0.05 with respect to 0μM Ac_5_GalNTGc. Ac_5_GalNTGc decreased mAb HECA-452 and CSLEX-1 binding, and upregulated Le^X^ expression.

### Ac_5_GalNTGc truncates O-glycan biosynthesis

A panel of lectins was applied to characterize changes in carbohydrate structures upon culture with Ac_5_GalNTGc (**Fig. 3, Supplemental Fig. S2**). Studies were performed both in the presence or absence of sialidase, as some lectins preferentially bind de-sialylated epitopes. Here, Ac_5_GalNTGc augmented VVA binding by 30-fold compared to vehicle treatment (Fig. 3A). Similar observations were also made with another GalNAcα binding lectin, Soyabean Agglutinin SBA (Fig. S2). In agreement with this, pronounced VVA binding to cells was observed in the fluorescent micrographs upon culture with Ac_5_GalNTGc (Fig. 3F). Compared to the sialidase treated [O]^−^ cells (COSMC-knock out) which augmented VVA binding by 43-fold, Ac_5_GalNTGc was 13-fold effective (Fig. 3A). Thus, Ac_5_GalNTGc truncates 1/3rd of the O-glycans at the Tn-antigen stage. Similar observations were also made in lectin blots, where Ac_5_GalNTGc dramatically elevated VVA binding to a variety of glycoproteins (Supplemental Fig. S3A). Besides leukocytic cells, a similar increase in VVA-binding was also noted on other breast (T47D, ZR-75-1) and prostate (PC-3) cell lines, but not HEK293T kidney cells (Supplemental Fig. S3B). Consistent with the notion that *O*-glycans are specifically altered, a 30% reduction in the sialylated T-antigen was observed in HL-60s, upon using PNA (Fig. 3B, Fig. S2). In addition, Ac_5_GalNTGc decreased the *N*-acetyl-lactosamine structures reported by ECL by ~40% (Fig. 3C). α2,3-sialic acid measured using MAL-II (Fig.3D), complex N-glycans reported by PHA-L (Fig. 3E) and a panel of additional 15+ lectins that report on Man, Fuc, GlcNAc and Gal related epitopes (Fig. S2) remained largely unchanged. Overall, the data suggest a potent effect of Ac_5_GalNTGc on cell-surface *O*-glycans, with less effect on other types of glycoconjugates.

**Figure 3.**
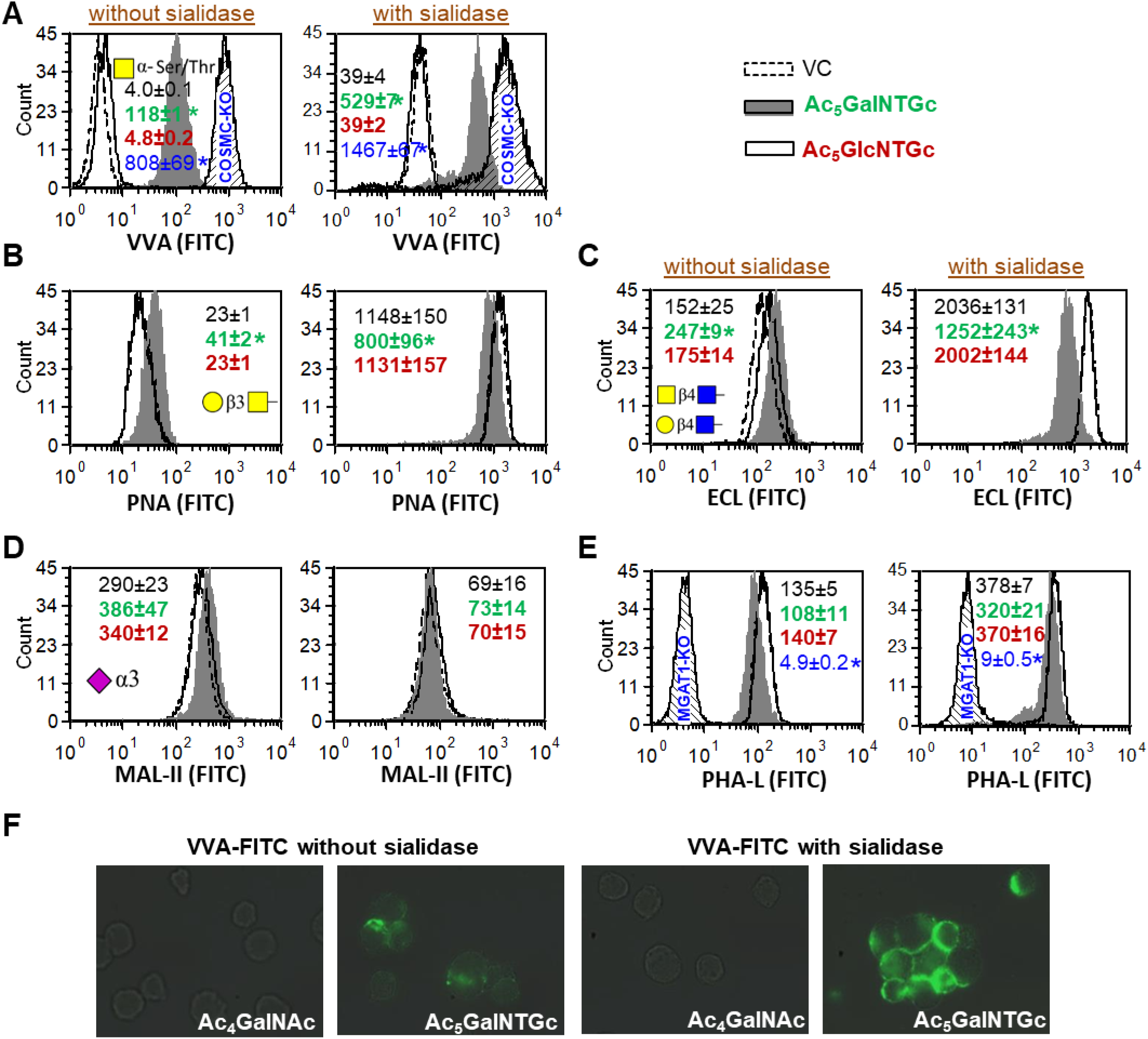
Lectin binding to GalNTGc treated HL60s. **A-F.** HL60s cultured with 80μM Ac_5_GalNTGc, Ac_5_GlcNTGc or VC for 40h were stained with labeled lectins, either before (left panels) or after (right panels) treatment with α2-3,6,8,9 *Arthrobacter Ureafaciens* neuraminidase: **A.** VVA (binds GalNAcα-Ser/Thr); **B.** PNA (Galβ1,3GalNAc); **C.** ECL (Galβ1,4GlcNAc); **D.** MAL-II (α(2,3)sialic acid); and **E.** PHA-L (complex N-glycans). Epitope bound by lectins is depicted in each panel using the Symbol Nomenclature For Glycans. Mean fluorescence intensity/MFI±SD are shown in individual panels for VC (top number, black), Ac_5_GalNTGc (second, green), and Ac_5_GlcNTGc (third, red). The fourth number (blue) depicts the MFI of CRISPR-Cas9 COSMC-KO cells that have truncated O-glycans (panel A) and MGAT1-KOs that have trucated N-glycans (panel E). * *P*<0.05 with respect to VC and Ac_5_GalNTGc. **F.** Fluorescence images of Ac_5_GalNTGc and Ac_4_GalNAc treated HL-60s, stained with VVA-FITC. Ac_5_GalNTGc increased VVA-lectin binding indicating the presence of truncated mucin-type O-glycans. Ac_5_GalNTGc reduced LacNAc and T-antigen formation without altering overall cellular sialylation.

### Ac_5_GalNTGc reduced leukocyte adhesion to L- and P-selectin

Microfluidics based flow chamber studies determined if the reduced sLe^X^ expression upon Ac_5_GalNTGc treatment attenuated leukocyte rolling on selectin-bearing substrates (**Fig. 4**). Here, Ac_5_GalNTGc treated HL-60s displayed 80% and 50% reduction in cell rolling on recombinant L-selectin (Fig. 4A) and on P-selectin bearing CHO-P cells (Fig. 4B), respectively. This reduction was not observed upon treatment with vehicle or GalNAc. In contrast to L- and P-selectin, Ac_5_GalNTGc did not alter E-selectin dependent leukocyte rolling on IL-1β stimulated HUVEC monolayers (Fig. 4C). P-selectin dependent leukocyte-platelet adhesion was also reduced by 40-50% upon cell culture with Ac_5_GalNTGc (Fig. 4D). Consistent with these functional data, Ac_5_GalNTGc reduced the apparent molecular mass of the major human L-/P-selectin ligand PSGL-1 by 15% (from 125 to 105KDa, Fig. 4E). It also reduced the mass of another common mucinous leukocyte glycoprotein, CD43. Together, the data demonstrate that O-glycan biosynthesis truncation by Ac_5_GalNTGc can reduce L- and P-selectin dependent leukocyte cell adhesion under shear, both in the context of leukocyte-endothelium and leukocyte-platelet binding.

**Figure 4.**
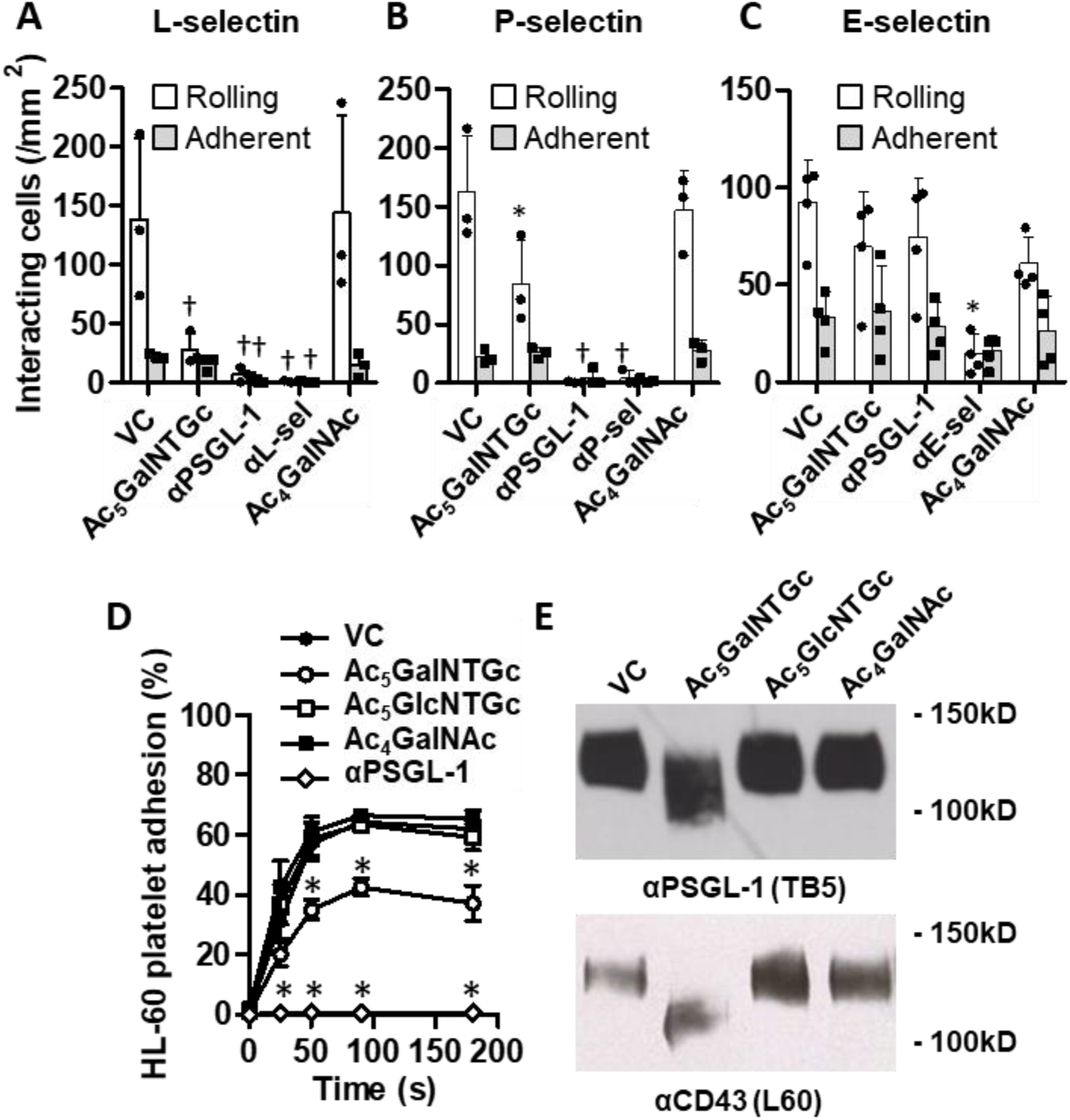
Effect of Ac_5_GalNTGc on cell adhesion. HL60 cells cultured with 80μM Ac_5_GalNTGc for 40h were perfused over substrates composed of: **A.** recombinant L-selectin, **B.** CHO-P cell monolayer bearing P-selectin, or **C.** IL-1β stimulated HUVEC monolayers bearing E-selectin. Wall shear stress was 1 dyne/cm^2^ in all cases. The density of rolling and adherent cells was quantified. Blocking antibodies used were against: PSGL-1 (αPSGL-1 mAb KPL-1), L-selectin (αL-sel DREG-56), P-selectin (αP-sel G1) and E-selectin (αE-sel HAE1f). **D.** HL60s were shear-mixed with TRAP-6 activated human platelets using a cone-plate viscometer at 650/s. Flow cytometry quantified % of HL-60s bound to at least one platelet. **E.** Western blots of HL60 cell lysates cultured with Ac_5_GalNTGc, Ac_5_GlcNTGc and Ac_4_GalNAc were probed with anti-human PSGL-1 antibody (mAb TB5) and anti-CD43 mAb L60. Ac_5_GalNTGc reduced cell adhesion to P- and L-selectin. It also reduced the molecular mass of PSGL-1 by truncating mucin-type O-glycans. * *P*<0.05 with respect to all other treatments, at indicated time. † *P*<0.05 with respect to other treatments excepts †’s are not different from each other.

### Ac_5_GalNTGc is pharmacologically active and it reduces granulocyte migration to sites of inflammation

To determine if Ac_5_GalNTGc is active *in vivo*, either mouse bone marrow cells (mBMCs) or neutrophils (mPMNs) were cultured with IL-3 (interleukin-3) and GCSF (granulocyte colony stimulating factor) *ex vivo* in the presence Ac_5_GalNTGc or control treatments for 40h (**Fig. 5A**). Similar to HL-60s, both Ac_5_GalNTGc treated mBMCs and mPMNs displayed a 6-fold increase in VVA binding at 40h (Fig. 5B). Ac_5_GalNTGc also reduced P-selectin-IgG binding to neutrophils by 50-65% (Fig. 5D) and L-selectin binding by 60-80% (Fig. 5E). PSGL-1 expression remained unchanged (Fig. 5C). In another set of studies, mBMCs cultured with Ac_5_GalNTGc or vehicle were labeled with distinct fluorescent dyes (either red or green), mixed in equal proportion, and injected *i.v.* into recipient mice in a thioglycollate-induced model of acute peritonitis. Twenty hours later, the peritoneal lavage was recovered, and the labeled neutrophils were enumerated using flow cytometry (Fig. 5F). Here, regardless of the labeling dye combination, Ac_5_GalNTGc reduced neutrophil migration into the peritoneum by ~50%, compared to vehicle control.

**Figure 5.**
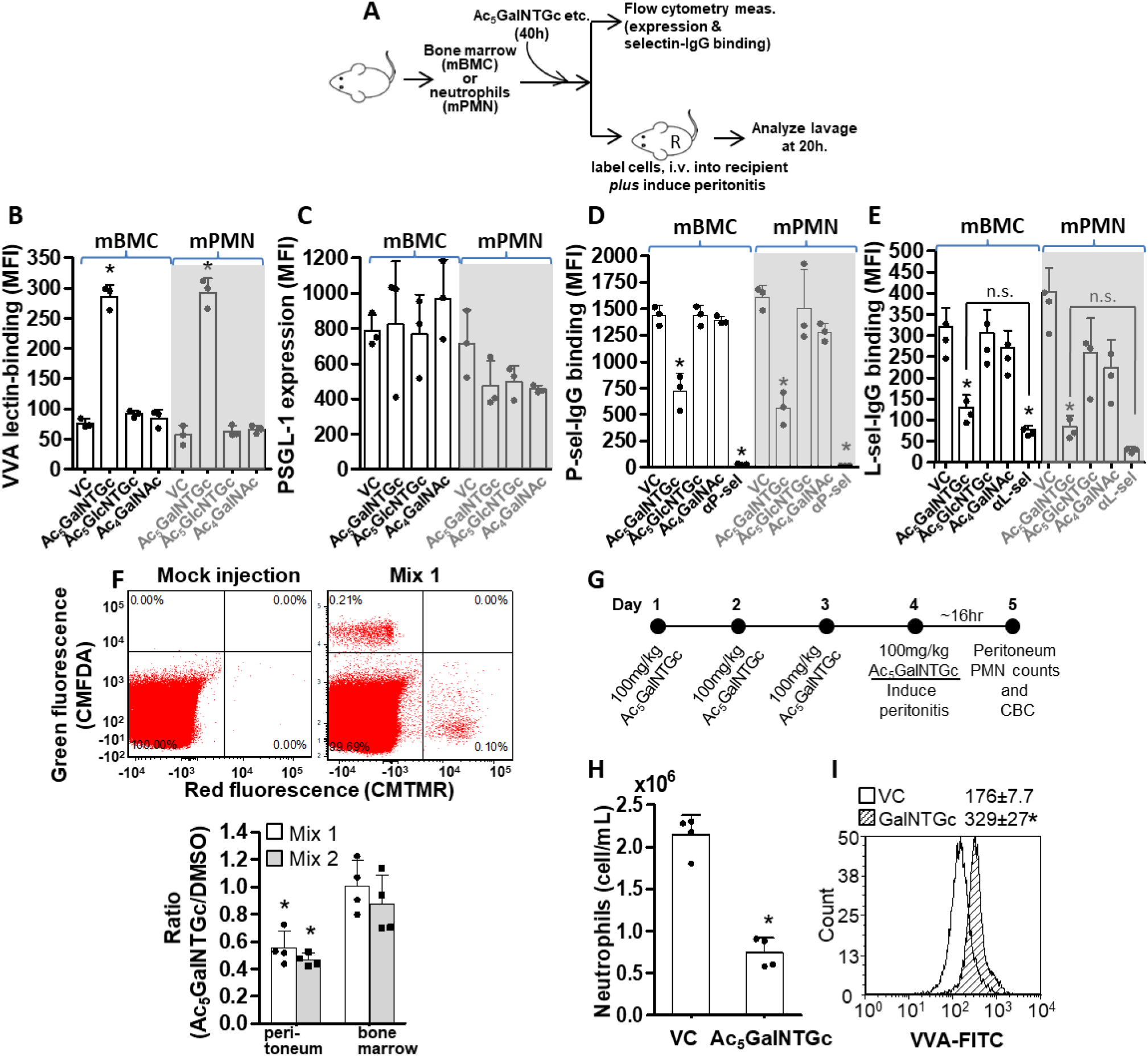
Murine acute inflammation. **A.** Mouse bone marrow cells (mBMC) and neutrophils (mPMN) were isolated from 10-12 week C57BL/6 mice, and cultured with 50μM HexNAc analogs or controls for 40h. Cells were analyzed using flow cytometry and used in a murine acute inflammation model. **B.** -**E.** Flow cytometry measured the binding of the following reagents to mouse neutrophils (CD11b+, Gr-1/Ly-6G/1A8+, F4/80-): **B.** VVA-FITC, **C.** Anti-mouse PSGL-1 mAb 2PH1, **D.** P-selectin-IgG and **E.** L-selectin-IgG. Ac_5_GalNTGc increased VVA-lectin binding by 4-5 fold, and reduced L-/P-selectin IgG binding by 50-70% without affecting PSGL-1 expression. **F.** mBMCs cultured with 80μM Ac_5_GalNTGc for 40h were mixed with VC at 1:1 ratio. In mix 1, Ac_5_GalNTGc cells were labeled with CMTMR (Red) while VC was CMFDA (Green) labeled. Labels were swapped in Mix 2 (dot plot not shown). Mix 1 or 2 cells were tail-vein injected into recipient mice following thioglycollate injection i.p. Red:green ratio of Gr-1+ cells in the peritoneal lavage and bone marrow was measured at 20h. Ac_5_GalNTGc reduced neutrophil counts in peritoneum by 50% in both Mixes. **G-I.** Ac_5_GalNTGc (100mg/kg/day) or VC was injected daily into mice for 4 days prior to induction of peritonitis. Murine neutrophil (CD11b+, Gr-1/Ly-6G/1A8+, F4/80-) counts in the peritoneum were quantified at 16h. Neutrophil counts in peritoneal lavage was reduced by 65% in Ac_5_GalNTGc treated mice (panel **H**). VVA binding was augmented in peritoneal neutrophils (panel **I**). \**P*<0.05 with respect to all other treatments in each panel, except as indicated.

Next, to confirm the pharmacological activity of Ac_5_GalNTGc, 100mg/kg Ac_5_GalNTGc was infused into recipient mice once daily for 4-days before induction of peritonitis (Fig. 5G). Here, also, Ac_5_GalNTGc reduced neutrophil extravasation into the peritoneal lavage by 65% (Fig. 5H). Peripheral blood counts and leukocyte differentials remained unchanged at day-4 (Table S1), and animals did not exhibit any signs of distress or abnormality. Significantly, the VVA binding to neutrophils in the peritoneal lavage was two-fold higher upon Ac_5_GalNTGc treatment (Fig, 5I). Together, the data confirm the *in vivo* metabolic activity of Ac_5_GalNTGc.

### Ac_5_GalNTGc blunts T-antigen biosynthesis, with less effect on N-glycans and glycolipids

We verified the effect of Ac_5_GalNTGc on overall O-glycosylation by feeding peracetylated GalNAc-OBn to cells cultured with Ac_5_GalNTGc or GalNAc (control), and, measuring extended glycans formed on this substrate (**Fig. 6A**). All products formed were verified based on MS/MS and also LC retention time. [NOG]^−^ HL-60s fed with GalNAc-OBn served as negative controls, since they lack the O-glycan biosynthesis machinery. Here, whereas 35% of the GalNAc-O-Bn was converted into extended O-glycans (disaccharides, trisaccharides etc.) in the GalNAc fed cells, this was reduced to 14% upon culture with Ac_5_GalNTGc and such biosynthesis was absent on [NOG]^−^ cells. These data suggest ~60% (=21/35×100) inhibition of O-glycan elaboration upon Ac_5_GalNTGc treatment. The increased prevalence of exposed unmodified GalNAc upon culture with Ac_5_GalNTGc is consistent with the increased VVA binding noted previously (Fig. 3). Due to reduced GalNAc-O-Bn substrate extension, longer glycan chains including those containing the T-antigen (Galβ1-3GalNAc/core-1) and core-2 glycan (Galβ1-3[GlcNAc β1-6]GalNAc) related structures were reduced in the Ac_5_GalNTGc treated cells. Such reduction in O-glycan extension could account for the reduced molecular mass of PSGL-1 and CD43. Notably, consistent with cytometry measurements, sLe^X^ epitope biosynthesis on core-2 O-glycans was reduced by ~70% in the Ac_5_GalNTGc treated cells. The direct incorporation of GalNTGc/HexNTGc (in non-acetylated or partially acetylated form) into carbohydrate products formed on the GalNAc-OBn substrate was not observed. In enzymology studies, a ~40% reduction in β1,3GalT activity was noted in the HL60s upon culture with Ac_5_GalNTGc, and this may partially explain the increased expression of the Tn-antigen epitope (Fig. 6B).

**Figure 6.**
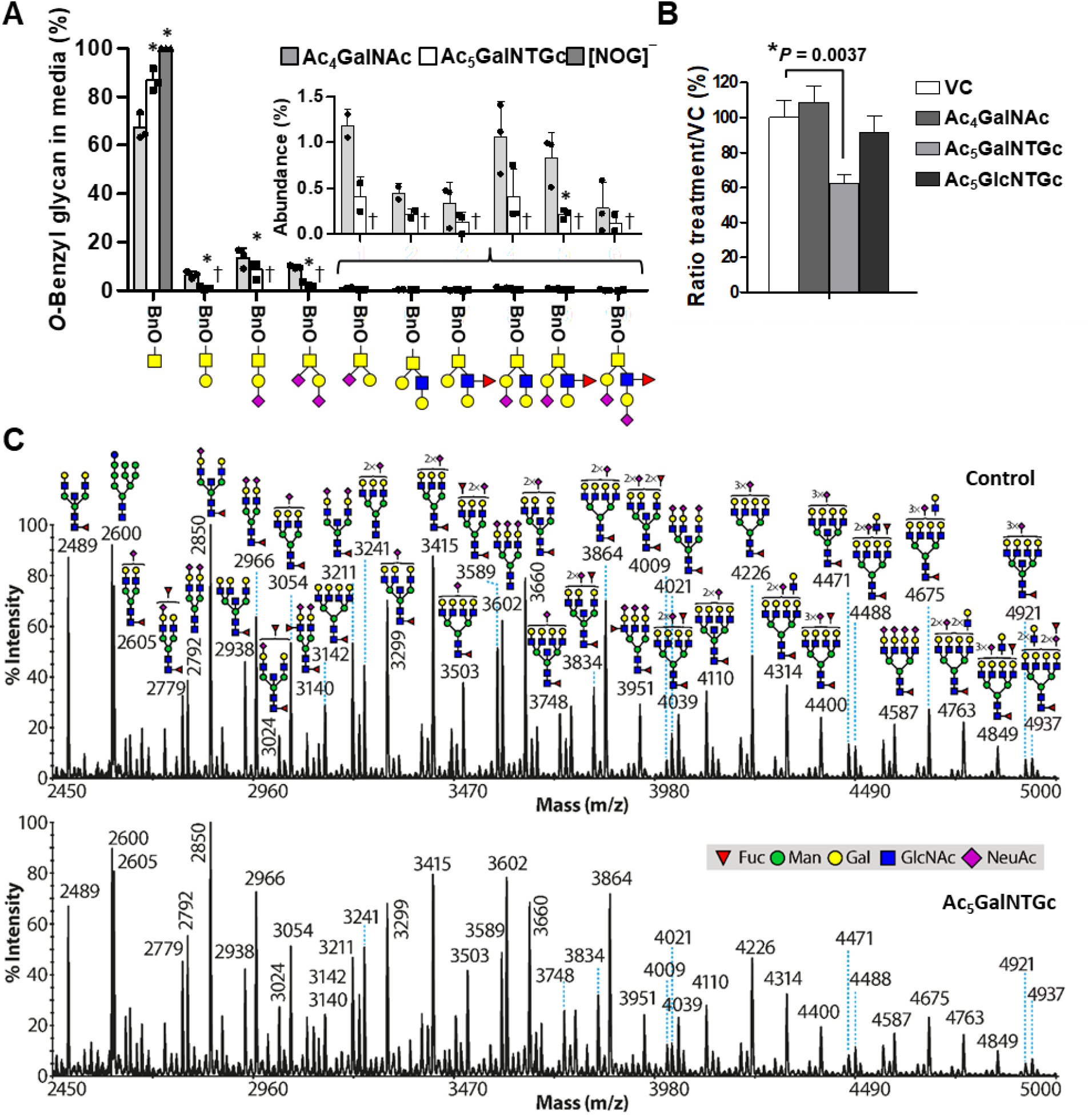
Effect of Ac_5_GalNTGc on O- and N-glycan biosynthesis. **A.** HL-60s were cultured with 80μM Ac_5_GalNTGc or Ac_4_GalNAc for 16h prior to addition of peracetylated GalNAc-O-Bn for an additional 48h. [NOG]^−^ TKO HL-60s cultured with peracetylated GalNAc-O-Bn served as negative control as they lack C1GalT1-chaperone COSMC activity. GalNAc-O-Bn and related products from cell culture supernatant were purified at the study end point, permethylated and analyzed using LC-MS/MS. Product ion-current area-under-the-curve quantified products formed. GalNAc-O-Bn consumption was reduced upon Ac_5_GalNTGc treatment and abolished in TKO HL60s. **B.** Cell lysates were prepared from HL60s cultured with 80μM Ac5GalNAc, Ac_5_GalNTGc or Ac_4_GalNAc for 40h. 3.3mg/mL lysate was mixed with 0.5mM GalNAc-O-Bn (substrate) and 1mM UDP-Gal donor overnight. β1,3GalT was quantified in LC-MS/MS runs by measuring ion current AUC (area under the curve) for product (Gal(β1-3)GalNAc-O-Bn) vs. unreacted substrate (GalNAc-O-Bn). Results are presented after normalization with respect to vehicle control (100%). Product was not formed in the absence of lysate (negative control). **C.** MALDI-TOF MS profile of permethylated N-glycans for cells treated with 80μM Ac_5_GalNTGc or vehicle for 40h. Putative structures are based on composition, tandem MS, and knowledge of biosynthetic pathways. All molecular ions are [M+Na]+. MALDI data are representative of duplicate runs. Ac_5_GalNTGc reduced T-antigen formation and O-glycan extension, without a major effect on N-glycan biosynthesis. * *P*<0.05 with respect to GalNAc treatment for each structure. † Glycan products were not detected.

MALDI-TOF MS based glycomics profiling was undertaken to study the impact of Ac_5_GalNTGc on cellular N-glycans and GSL biosynthesis. Here, Ac_5_GalNTGc did not have any significant impact on N-glycan structures, with high molecular weight complex structures still being observed (Fig. 6C). In the case of GSLs, however, Ac_5_GalNTGc appeared to quantitatively affect the abundance of certain GSLs mainly containing fucose residues (*m*/*z* 1566, 1841, 2016, 2639, 3088 and 3262, Supplemental Fig. S4). Direct incorporation of HexNTGc into glycoconjugates was not detected. Overall, the data suggest that Ac_5_GalNTGc reduces the formation of the T-antigen on leukocyte *O*-glycans and it also has some impact on GSLs. The effect of the compound on N-glycan biosynthesis was negligible.

### Minimal changes in sugar-nucleotide biosynthesis and low levels of Ac_5_GalNTGc derivative incorporation into cellular O-glycans, N-glycans and glycolipids

Recent studies suggest that monosaccharide analogs, where selected hydroxyl groups are modified with different substituents, often act as metabolic inhibitors by altering cellular sugar-nucleotide compositions (Del Solar et al., 2020; Gloster and Vocadlo, 2012; van Wijk et al., 2015). However, this is not the mechanism of action of Ac_5_GalNTGc since it did not alter cellular sugar-nucleotide levels based on LC-MS/MS (**Fig. 7A**). Here, glucose based sugar-nucleotide standards were distinguished from corresponding galactose counterparts based on retention time since compounds containing Glc eluted first, i.e. UDP-Glc eluted prior to UDP-Gal, and UDP-GlcNAc before UDP-GalNAc (Del Solar et al., 2020). Based on these observations, the MS/MS fragmentation pattern of UDP-HexNTGc (Supplemental Fig. S5A) and the observed elution chromatogram (Fig. S5B), our data suggest the possible conversion of Ac_5_GalNTGc into both UDP-GalNTGc and UDP-GlcNTGc in equal parts (Fig. 7B). While exact quantitation is not possible due to the absence of UDP-HexNTGc standards, MS ion counts of UDP-GalNTGc and UDP-GlcNTGc in cell lysates was 10-15 fold lower than that of UDP-GalNAc and UDP-GlcNAc suggesting only small amounts of UDP-HexNTGc formation.

**Figure 7.**
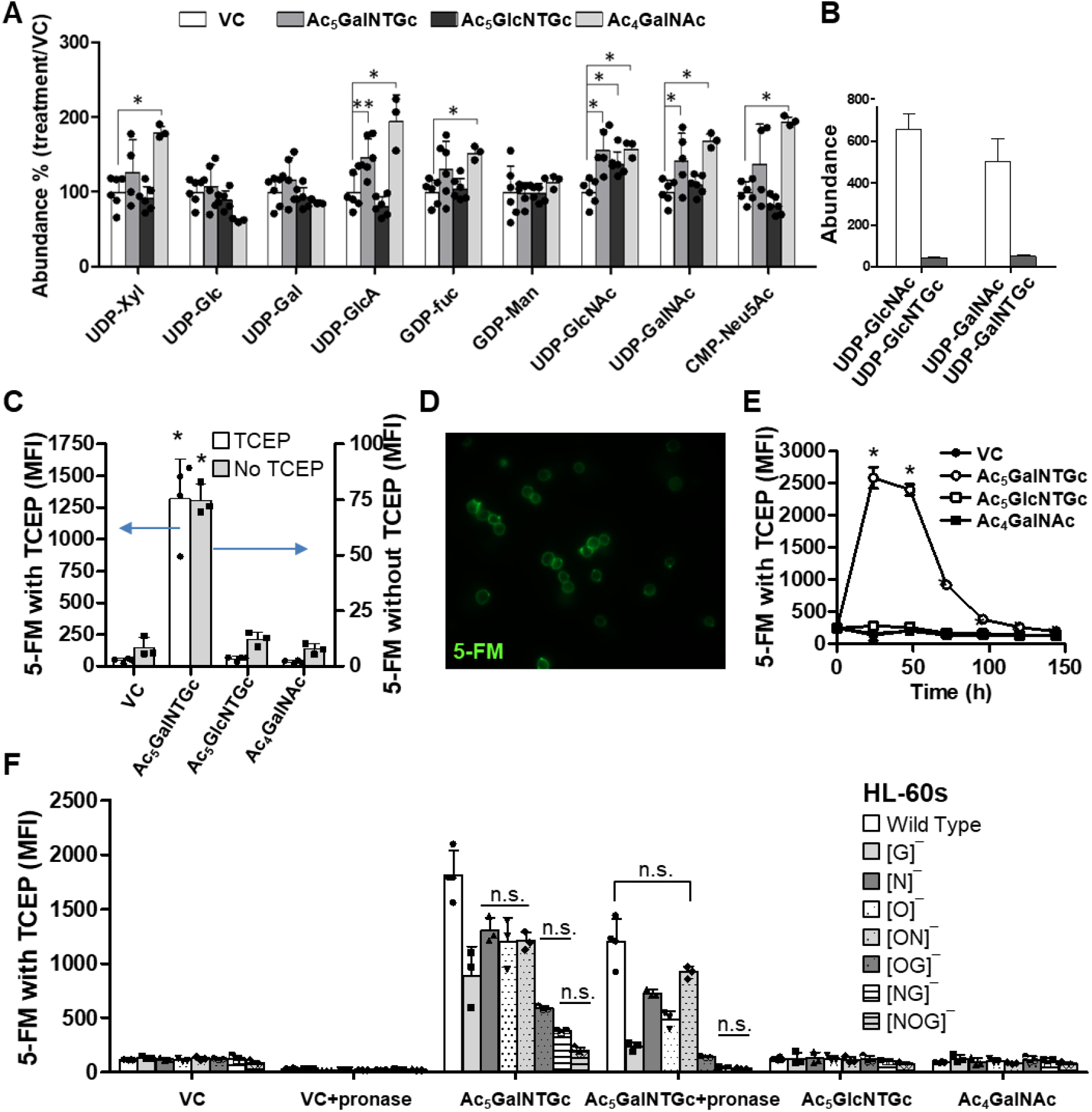
Direct incorporation of thiol into Ac_5_GalNTGc treated HL60 wild-type and knockout cells. HL60s were cultured with peracetylated HexNAc analogs or VC (80μM, 40-48h). **A.** Sugar nucleotide levels in cells determined using LC-MS/MS. Abundance are quantified based on area under MS^1^ curve for individual species, normalized with respect to vehicle control which is set to 100 (\**P*<0.05) **B.** UDP-HexNAc and UDP-HexNTGc abundance in GalNTGc (80μM, 48h) treated cells. Abundance are presented relative to CMP-Neu5Gc internal standard ionization in all runs. UDP-HexNTGc may exist in both UDP-GalNTGc and UDP-GlcNTGc forms. UDP-HexNTGc was not detected in vehicle, GalNAc and GlcNTGc samples. **C.** Cells were reacted with fluorescein-5-maleimide/5-FM, in the presence (left axis) or absence (right axis) of 10mM TCEP. A similar increase in 5-FM binding was noted in the GalNTGc treated cells under both conditions, although the signal was brighter following TCEP mediated reduction. **D.** Fluorescence microscopy showing the reaction of maleimide with free sulfhydryl groups predominantly on the cell surface. **E.** HexNAc-analogs incorporation time course. GalNTGc incorporation peaked 24-48h after compound addition. \**P*<0.05 with respect to other treatments. **F.** HexNAc analogs were cultured with wild-type HL-60s and a panel of isogenic CRISPR/Cas9 HL60-KO cell lines containing truncated O-glycans ([O]^−^), N-glycans ([N]^−^), GSLs ([G]^−^), dual knockouts ([ON]^−^, [OG]^−^, [NG]^−^) and triple knockouts ([NOG]^−^). 1 mg/ml pronase was added to some of these cells for 2 h to cleave cell-surface glycoproteins prior to reanalysis using the flow cytometer. 5-FM was incorporated into N-glycans, O-glycans and GSLs. All treatments in the GalNTGc samples were statistically different with respect to each other, except those indicated by ‘n.s.’ (not significant).

The direct incorporation of Ac_5_GalNTGc and its derivatives into glycoconjugates was not observed in MS studies (Fig. 6), potentially due to their low abundance, below the instrument detection limit. Low levels of GalNTGc incorporation was also observed in O- and N-glycans in LC-MS/MS glycoproteomics investigations (data not shown). However, this could be readily detected using fluorescence methods. Here, consistent with a previous report (Agarwal et al., 2013), we observed a 10-15 fold increase in FITC-maleimide (5-FM) incorporation upon culture with Ac_5_GalNTGc, but not in controls containing Ac_5_GlcNTGc or Ac5GalNAc (Fig. 7C). 5-FM incorporation was 15-fold higher when the labeling reaction was performed in the presence of the reducing agent TCEP, suggesting that a majority of the cell-surface thiol groups introduced by Ac_5_GalNTGc were crosslinked via disulfide bridges under native conditions, i.e. they exist as GalNTGc-GalNTGc or GalNTGc-Cys moieties. Similar results were obtained for cell surface thiol measurement by flow cytometry wherein the two-step maleimide-PEG2-biotin Michael addition was followed by FITC-conjugated avidin staining (Supplemental Fig. S6A). Cell surface incorporation of GalNTGc derivatives was also observed using fluorescence microscopy (Fig. 7D), but not in controls runs that used Ac_5_GlcNTGc or Ac5GalNAc (Fig. S6B). Ac_5_GalNTGc and its derivatives were maximally incorporated 24-48h post-treatment, with 5-FM signal being reduced to basal levels at 72-96h (Fig. 7E). Besides HL-60s, 5-FM was also incorporated into other human cell lines including breast (T47D, ZR-75-1), prostate (PC-3) and to a lesser degree into kidney (HEK293T) cells (Fig. S6C).

Studies were performed with a panel of CRISPR-Cas9 HL-60 knockouts that contain truncated O-glycans, N-glycans and/or GSLs, in order to determine the glycoconjugates that incorporate GalNTGc-derivatives (Fig. 7F). This analysis suggests the increased prevalence of sulfhydryl groups in all families of cell-surface glycans, with incorporation being somewhat higher in GSLs (~40-50%) compared to N-glycans (~25-35%), followed by O-glycans (~20-25%). These estimates are based on the quantitative incorporation of 5-FM signal in the single and double knockout cell lines compared to wild-type HL60s. 5-FM signal was low/absent in the triple knockouts and thus this signal comes from thiol incorporation into glycans and not into other macromolecules. It could also be reduced upon protease digesting cell surface glycoproteins. Consistent with the above, maleimide incorporation was observed in western blots of PSGL-1 with greater maleimide incorporation being noted for lower molecular mass glycoproteins that contain more truncated O-glycans (Fig. S7A). Additionally, Ac_5_GalNTGc may not be transformed into sialic acid using the pathway illustrated in Fig. S7B, since the removal of sialic acids by α2-3,6,8,9-neuraminidase did not reduce the measured 5-FM signal. Overall, the data are consistent with the notion that a portion of Ac_5_GalNTGc is converted into UDP-derivatives that are incorporated into cellular glycoproteins and possibly also GSLs.

## DISCUSSION

Our results demonstrate that Ac_5_GalNTGc is a potent metabolic inhibitor of O-glycan biosynthesis in diverse cell types including mouse peripheral blood neutrophils, human promyelocytes, breast and prostate cancer cell lines. In all these cells, Ac_5_GalNTGc upregulated VVA binding suggesting that it may reduce core1 β1,3GalT1 (C1GalT1) activity. Upon quantitatively comparing VVA binding on Ac_5_GalNTGc treated cells with COSMC-knockouts that completely lack C1GalT1 activity, it is estimated that at least ~1/3^rd^ of the HL-60 O-glycans are truncated by Ac_5_GalNTGc as GalNAc bearing polypeptides with no extension. This conclusion is consistent with the ~20-30% decrease in molecular mass of mucinous proteins including PSGL-1 and CD43 on the leukocytes. Additionally, feeding peracetylated GalNTGc to these cells reduced Gal incorporation into benzyl-GalNAc by ~60% with respect to GalNAc control, with few extended O-glycan structures. N-glycan profiling using MALDI TOF MS confirmed minimal effect of Ac_5_GalNTGc on N-glycan biosynthesis. The compound, however, appeared to affect the abundance of some fucosylated GSLs via yet unidentified mechanisms. Finally, although, the current study did not examine glycosaminoglycan and O-GlcNAc type modifications, the results thus far suggest that the major impact of Ac_5_GalNTGc is on O-glycan biosynthesis with the compound reducing the elaboration of such entities by ~30-60%. Additionally, Ac_5_GalNTGc has greater potency compared to peracetylated GalNAc-O-Bn, a previously described O-glycan inhibitor which is added at 2-4mM into cell culture medium to reduce O-glycan biosynthesis (Alfalah et al., 1999; Huet et al., 1998; Kuan et al., 1989; Tsuiji et al., 2003). In contrast, the functional effect of Ac_5_GalNTGc was observed at concentrations as low as 10μM with maximum efficacy at 50-80μM. The lower usage dose and favorable pharmacological properties allow systemic usage of Ac_5_GalNTGc in murine models.

The functional effects of Ac_5_GalNTGc on O-glycan biosynthesis inhibition was assessed in a model of inflammation where leukocytes were recruited onto selectin-bearing substrates under shear flow. Here, Ac_5_GalNTGc dramatically reduced cell surface sLe^X^ expression as measured using mAbs HECA-452 and CSLEX-1, and also using O-glycan mass spectrometry analysis. Such inhibition was prominent on mucinous proteins like the P- and L-selectin ligand PSGL-1, as the apparent mass of this glycoprotein was reduced by ~25% upon culture with Ac_5_GalNTGc. Ac_5_GalNTGc treatment also reduced leukocyte adhesion on L- and P-selectin, but not E-selectin, under hydrodynamic shear. This is consistent with a previous report that C1GalT1 in necessary for leukocytes recruitment via L- and P-selectin (Stolfa et al., 2016). Abolishing O-glycosylation, however, has only a minor effect on human leukocyte recruitment/tethering and rolling on E-selectin. Besides the effect on cell rolling, Ac_5_GalNTGc also reduced the extent of P-selectin-PSGL-1 dependent leukocyte-platelet adhesion under shear. Mouse neutrophils (and mBMCs) cultured *ex vivo* with Ac_5_GalNTGc for 40h also exhibited higher than normal levels of VVA binding, and reduced interaction with L- and P-selectin IgG fusion proteins. In the complex *in vivo* milieu during peritonitis, consistent with the inhibition effect of Ac_5_GalNTGc on L-/P-selectin dependent binding, neutrophil recruitment to sites of inflammation was reduced upon culture of cells with Ac_5_GalNTGc. Ac_5_GalNTGc also had excellent pharmacological properties, and it caused 60% reduction in leukocyte recruitment at sites of inflammation. These data support the use of Ac_5_GalNTGc *in vivo* for anti-inflammatory therapy.

Studies were undertaken to determine the mechanism of Ac_5_GalNTGc action, with focus on β1,3GalT activity, since addition of this compound into cell culture medium both increased VVA binding and drastically reduced Gal incorporation into GalNAc-O-Bn substrate. β1,3GalT enzymatic activity was also partially reduced upon culture with Ac_5_GalNTGc, consistent with the notion that this compound may reduce core-1 glycan biosynthesis. In such investigations, we did not observe marked changes in the cellular nucleotide-sugar profile of HL60s culture with Ac_5_GalNTGc, that would indicate non-specific activity of the compound. This is unlike previous studies that used modified monosaccharides, which globally alter the cellular nucleotide-sugar profile (Del Solar et al., 2020; Rillahan et al., 2012; van Wijk et al., 2015). Here, culture of cells with per-acetylated 6F-GalNAc reduced the cellular UDP-GalNAc and UDP-GlcNAc pool by ~80-90% (van Wijk et al., 2015), 2F-Fuc also depressed cellular GDP-Fuc, and 3F-Neu5Ac similarly abolished CMP-Neu5Ac (Rillahan et al., 2012). In these studies, substantial amounts of UDP-(6F)GalNAc, GDP-(2F)Fuc and CMP-(3F)NeuAc were formed and this resulted in collateral reduction in corresponding unmodified nucleotide-sugars. Unlike this, the culturing of cells with Ac_5_GalNTGc resulted in relatively low levels of UDP-HexNTGc synthesis. Based on ion count data, both UDP-GalNTGc and UDP-GlcNTGc were formed via the salvage pathway, although the levels were 1/10-1/15th that of UDP-GalNAc and UDP-GlcNAc. Due to this, only low levels of GalNTGc and GlcNTGc were integrated into cellular glycolipids, N-glycans and O-glycans. This could be detected using sensitive fluorescence based methods (flow cytometry and microscopy), but not mass spectrometry. In this regard, the degree of GalNTGc integration into glycoproteins may be important for functional efficacy since cell systems with greater maleimide-FITC incorporation (e.g. HL-60, T47D, PC3) also displayed greater enhancement of VVA binding. HEK cells, on the other hand, exhibiting low maleimide incorporation and minimal change in VVA engagement. Transcriptional analysis of these cells does not reveal any obvious differences in O-glycosylation related enzyme expression profiles to explain these observations (Supplemental Table S2), though nucleotide-sugar analysis needs to be performed to quantify the efficiency of UDP-HexNTGc synthesis across these different cell types. Additionally, if Ac_5_GalNTGc reduces T-synthase activity, one possibility is that glycoproteins containing directly incorporated GalNTGc may bond directly with either C1GalT1 or its unique molecular chaperone COSMC, within the Golgi (Ju et al., 2002; Ju and Cummings, 2002). Such molecular interactions may occur between the thiol residues of GalNTGc-derivatives and free Cys available on C1GalT1/COSMC. These may reduce T-synthase activity. In this regard, previous studies show that both C1GalT1 and COSMC are highly conserved proteins with 363 and 318 amino acid residues, respectively (Ju et al., 2002; Ju and Cummings, 2002). They both contain 6 highly conserved Cys residues in their luminal/catalytic domain including a pair of vicinal Cys that may be targeted by GalNTGc containing glycoproteins. Alternatively, the steric repulsion of the bulky thiol groups may preclude binding of the GalNTGc-decorated polypeptide acceptors to C1GalT and resulting enzyme activity. Other factors that may contribute to Ac_5_GalNTGc inhibitory function include: i. potential roles for partially deacetylated GalNTGc or its derivative in regulating glycosylation; ii. inhibition of other enzyme activities besides C1GalT1/COSMC, particularly those related to ppGalNAcT function. Additional studies are needed to examine these hypothesis.

Overall, Ac_5_GalNTGc is a potent inhibitor of O-linked glycosylation. It fills an important gap in the field that lacks O-glycosylation inhibitors. The compound is pharmacologically active, it reduces the expression of sialofucosylated glycan epitopes on the leukocyte cell surface, inhibits L- and P-selectin-dependent molecular recognition under static and flow conditions, and displays the potential to have anti-inflammatory properties. Besides basic science applications, the compound may find utility in translational studies where there is a need to trim mucinous glycoproteins for example during pulmonary disorders with excess mucin production, and in the context of investigations related to cancer metastasis and immunotherapy.

## SIGNIFICANCE

Four common types of glycans are expressed on the surface of mammalian cells. These include *O*- and *N*-linked glycans on glycoproteins, glycosphingolipids (GSLs) and glycosaminoglycans (GAGs). Currently, there are a number of ways to study *N*-glycan function using enzymes like PNGaseF/Peptide:N-glycosidase F that cleave these structures, and glycosidase inhibitors (e.g. kifunensine) that can truncate *N*-glycan biosynthesis. Small molecule inhibitors also exist for the study of GSLs (e.g. D-threo-1-phenyl-2-palmitoylamino-3-morpholino-l-propanol/PPMP), and lyases are commonly used to trim GAGs. Unlike these, few reagents are available to study mucin type *O*-glycosylation. The most common chemical method to block *O*-glycan biosynthesis involves the use of a small molecule called peracetylated benzyl-GalNAc. This compound is however only effective when used at millimolar concentrations and this limits *in vivo* usage. To address the above limitations regarding the need for new tools to study *O*-linked glycosylation, this manuscript presents the characterization of a peracetylated *N*-thioglycolyl modified GalNAc analog (‘Ac_5_GalNTGc). Our data show that Ac_5_GalNTGc is an efficient inhibitor of *O*-glycan biosynthesis at concentrations as low as 50 μM. At these doses, Ac_5_GalNTGc abolished sialyl Lewis-X (sLeX) epitope expression on human leukocyte *O*-glycans, and inhibited leukocyte rolling on L- and P-selectin substrates under flow. Addition of this compound to diverse cell types resulted in a dramatic upregulation of VVA-lectin binding, suggesting that Ac_5_GalNTGc acts by inhibiting core-1 *O*-glycan elaboration. Ac_5_GalNTGc did not alter cellular nucleotide-sugar levels or *N*-glycan biosynthesis. Thus, it primarily inhibits *O*-linked glycosylation. The compound was also pharmacologically active in mouse, and it inhibited neutrophil recruitment to sites of inflammation in a model of acute peritonitis. Ac_5_GalNTGc may find diverse *in vitro* and *in vivo* applications as a potent inhibitor of O-linked glycosylation.

## Acknowledgements

Supported by the NIH awards HL103411 and GM126537 to SN, and the Biotechnology and Biological Sciences Research Council Grant BB/K016164/1 to AD and SMH. V.dS was partially supported by the University at Buffalo Genome, Environment and Microbiome program.

## Author contributions

Conceptualization, S-G. S., S.N.; Writing-Original Draft: S.S.W., S.N.; Writing-Review and Editing: all authors; Supervision: J.T.L., A.D., S.M.H., S-G. S., S.N.; Investigation and Methodology: S.S.W., V.dS, X.Y., A.A., A.E.F., M.N., A.A.; Validation: V.dS, K.A., M.G., S.M.A.; Resources: M.G., S-G.S.; Funding Acquisition: A.D., S.M.H., S.N.; Co-Corresponding authors: S-G. S. lead the chemical synthesis effort and some of the *in vitro* studies using HL-60s. S.N. lead other functional studies.

## Declaration of interests

We do not have competing financial or non-financial interests.

## STAR METHODS

### LEAD CONTACT AND MATERIALS AVAILABILITY

Further information and requests for resources and reagents should be directed to and will be fulfilled by the Lead Contact, Sriram Neelamegham.

## EXPERIMENTAL MODEL AND SUBJECT DETAILS

8-12 week-old C57BL/6 wild-type mice of either sex were used. Animals were randomized prior to experimentation. All animal studies were approved by the Roswell Park Cancer Institute Animal Care and Use Committee (RPCI-IACUC). HL-60 cells (female promyeloblasts, RRID:CVCL_0002), T47D (female ductal carcinoma, RRID: CVCL_0553) and ZR-75-1 (female epithelial ductal carcinoma, RRID: CVCL_0588), metastatic prostate PC-3 cells (male adenocarcinoma, RRID: CVCL_0035) and embryonic kidney HEK293T (fetal epithelial, RRID:CVCL_0063) were obtained from ATCC. Human Umbilical Vein Endothelial Cells (HUVECs, cat #CC-2519A) were from Lonza.

## METHODS DETAILS

### Reagents

The synthesis of peracetylated 1,3,4,6-tetra-*O*-acetyl-2-acetamido-2-deoxy-α-D-galactopyranose (abbreviated ‘Ac_4_GalNAc’), 1,3,4,6-tetra-*O*-acetyl-2-(2-acetylthio)acetamido-2-deoxy-β-D-galactopyranose (Ac_5_GalNTGc), 1,3,4,6-tetra-*O*-acetyl-2-acetoxyacetamido-2-deoxy-β-D-galactopyranose (Ac5GalNGc), 1,3,4,6-tetra-*O*-acetyl-2-deoxy-2-propanamido-β-D-galactopyranose (Ac4GalNPr) and 1,3,4,6-tetra-*O*-acetyl-2-azidoacetamido-2-deoxy-β-D-galactopyranose (Ac4GalNAz) were previously described (Agarwal et al., 2013) (Fig. 1). All other chemicals were from ThermoFisher or Sigma.

All antibodies were mouse IgGs from BD Biosciences (San Jose, CA) unless otherwise mentioned. These include anti-CD15/Lewis-X mAb HI98 (IgM), rat anti-Cutaneous Lymphocyte Antigen mAb HECA-452 (IgM), anti-CD15s/sLe^X^ mAb CSLEX1, anti-CD162/PSGL-1 mAbs KPL-1 and TB5 (GeneTex, Irvine, CA), anti-CD43 clone L60, and isotype controls. The anti-CD65s mAb VIM-2 (IgM) was from AbD Serotec (Oxford, UK). Among these, both mAbs HECA-452 and CSLEX-1 bind overlapping sLeX and related sialylofucosylated glycans. Mouse cells were stained with either FITC rat anti-mouse Ly-6G and Ly-6C clone RB6-8C5, PE/APC rat anti-mouse Ly-6G mAb 1A8 (BioLegend, San Diego, CA), or PE rat anti-mouse CD162 mAb 2PH1. Function blocking mAbs that block selectin-mediated binding include anti-CD62P/P-selectin mAb G1, anti-CD62E/E-selectin mAb HAE-1f (Ancell, Bayport, MN) or P2H3 (eBioscience, San Diego, CA), and anti-CD62L/L-selectin mAb DREG-56. All lectins were purchased from Vector Laboratories (Burlingame, CA) as unlabeled or fluorescein-conjugated products. Supplemental Material provides lectin specificity data. In some instances, these reagents were labeled *in house* using Alexa-488 dye as described elsewhere (Yang et al., 2020).

### Cell culture and inhibitor treatment

Human promyelocytic leukemia cells (HL60s) were cultured in Iscove’s Modified Dulbecco’s Medium (IMDM) with 10% FBS. All other tumor cell lines were from ATCC (Manassas, VA) and cultured according to supplier’s instructions. CHO-S (Chinese Hamster Ovary) cells stably expressing P-selectin (CHO-P) were maintained in Dulbecco’s Modified Eagle’s Medium (DMEM) containing 10% FBS. Human umbilical vein endothelial cells (HUVECs) were cultured in EBM-2 basal medium (Lonza, Walkersville, MD). A panel of seven CRISPR-Cas9 knockout isogenic HL-60 clones displaying truncated glycoconjugates were available from a previous study (Stolfa et al., 2016). These include: i. [O]^−^ cells, which have truncated O-linked glycans due to genomic deletion of the core-1 β1,3galactosyltransferase chaperone Cosmc; ii. [N]^−^ cells that do not contain hybrid and complex N-glycans due to genomic deletion of Mgat1 (α-1,3-mannosyl-glycoprotein β1,2-N-acetylglucosaminyltransferase) activity; iii. [G]^−^ cells without GlcCer glycolipids due to absence of UGCG (UDP-glucose ceramide glucosyltransferase); iv-vi. dual-KOs lacking COSMC and Mgat1 ([ON]^−^ HL-60s), COSMC/UGCG-KOs ([OG]^−^ HL-60s) and Mgat1/UGCG-KOs ([NG]^−^ HL-60s); and vii. a triple knockout lacking all three glycoconjugates (Mgat1/COSMC/UGCG triple TKOs or [NOG]^−^ HL-60s). Absence of specific enzyme activity in these KOs was confirmed using Sanger sequencing, enzymology assays and lectin-binding based flow cytometry (Stolfa et al., 2016).

Peracetylated HexNAc compounds were dissolved in anhydrous dimethyl sulfoxide (DMSO) at 40mM stock. During experimentation, 0-200μM of these analogs were diluted into culture media containing 0.5-1×10^6^ cells/ml. Functional studies were performed at specified time points.

### Flow cytometry

Flow cytometry analysis was performed using either a BD FACSCalibur or Fortessa X-20 cytometer. Here, 2×10^6^ cells/ml suspended in HEPES buffer (30 mM HEPES, 110 mM NaCl, 10 mM KCl, 2 mM MgCl_2_, 10 mM glucose, 1.5mM CaCl_2_ containing 0.1% human serum albumin/HSA, pH 7.3) were labeled using 1-10μg/ml antibodies/lectins for 20 min at 4°C. In some runs, terminal sialic acid was removed by adding 0.1 U/ml α2-3,6,8,9-neuraminidase from *Arthrobacter ureafaciens* (New England Biolabs, Ipswich, MA) for 1h at 37°C. In other runs, cells were incubated with or without 10mM TCEP (Tris(2-carboxyethyl)phosphine) at pH 7.3 for 5-10 min, before 2μM Fluorescein-5-Maleimide (5-FM, Thermo-Pierce) addition at 4°C for 1h. Additionally, 1mg/ml of pronase (Roche, Indianapolis, IN) was sometimes added to cells for 2h at 37°C to remove surface proteins. Following labeling, the cells were washed, and resuspended in fresh HEPES buffer for cytometry analysis.

### Fluorescence microscopy

HL60 cells, with or without GalNAc analog treatment, were labeled with fluorescent VVA lectin as in the above cytometry studies. In some cases, the cells were α2-3,6,8,9-neuraminidase treated prior to VVA-labeling. Following fixation using 0.5% paraformaldehyde, the cells were mounted using Prolong Gold Antifade Reagent (Invitrogen) following manufacturer’s instructions. Phase contrast and fluorescence images were acquired using a Zeiss AxioObserver Z1 microscope (Plan-Apochromat 63x/1.40 Oil DIC M27 objective).

### SDS-PAGE and Western Blot

Cells were lysed using RIPA buffer containing HaltTM protease inhibitor (Thermo-Pierce) for 30-45 min on ice. Following centrifugation at 14,000g, the supernatant was collected and boiled in Laemmli sample buffer containing β-mercaptoethanol. Lysates from ~0.3-1×10^6^ cells were loaded in each well of either a standard 7.5% or 4-20% gradient gel (Thermo-Fisher). Following SDS-PAGE and transfer onto nitrocellulose, the membranes were probed using either mAb TB5, HECA-452 or L60 (1:1000 dilution), followed by 1:2500 HRP conjugated secondary Ab (Jackson Immuno, West Grove, PA) and enhanced chemiluminescence detection.

In some runs, the cells were treated with 10mM TCEP for 5-10 min. at room temperature, and then free-sulfhydryl groups were labeled using EZ-Link® Maleimide-PEG2-Biotin (1mM, 2h, 4°C, Thermo-Fisher) (Sampathkumar et al., 2006). The cells were then washed, lysed as above and PSGL-1 was immunoprecipitated using protein A/G agarose beads. Western blotting was performed using either anti-PSGL-1 mAb TB5 or HRP-linked anti-biotin (Cell Signaling, Danvers, MA).

For lectin blots, 20 μg of total cell lysate prepared as above were resolved using 7.5% SDS-gel and transferred onto nitrocellulose. Membranes were then blocked using 3.0 % gelatin in PBS-tween-20 (0.1%) for 1h at RT, incubated with 1:2000 biotinylated VVA lectin in PBS-tween-20 (0.1%) for 1h at RT, followed by 1:25,000 HRP conjugated avidin in 1% BSA in PBS-tween-20 (0.1%) for 1h at RT. Blots were developed using ECL substrate. Silver staining was performed in parallel using gels loaded with 2.0 μg cell lysate/lane.

### Microfluidic flow chamber based cell adhesion assay

A custom flow chamber with dimensions of 0.4mm(W) × 0.1mm(H) × 1cm(L) was fabricated using polydimethylsiloxane (Buffone et al., 2013). This was vacuum-sealed on a tissue culture plastic Petri dish and mounted on the stage of an inverted Zeiss AxioObserver Z1 microscope. The flow chamber substrate was composed of CHO-P cell monolayer expressing P-selectin, IL-1β stimulated HUVEC monolayers expressing E-selectin, or L-selectin-Fc fusion protein that was incubated overnight at 25 μg/mL and subsequently blocked with 1% bovine serum albumin (BSA). 2×10^6^ HL-60s/mL suspended in HEPES buffer were perfused over these substrates at 1 dyn/cm^2^. Movies of the cell interactions were recorded using a pco.edge sCMOS camera (Kelheim, Germany) and data were analyzed as described previously (Mondal et al., 2015).

### HL-60-platelet adhesion

Blood was collected from healthy human adult volunteers into 1:9 sodium citrate by venipuncture, following protocols approved by the University at Buffalo Institutional Review Board. Platelet rich plasma (PRP) was isolated and labeled using BCECF (2’,7’-Bis-(2-Carboxyethyl)-5-(-and-6)-Carboxyfluorescein, acetoxymethyl ester) (Zhang et al., 2018). 1:10 diluted PRP, 10μM TRAP-6 (Thrombin receptor activating hexapeptide) and 1×10^6^ HL-60s/mL were then shear mixed at 650/s in a cone-plate viscometer (VT-550, Thermo-Haake). Samples collected at various times were analyzed using a FACSCalibur cytometer to quantify % platelet-HL-60 binding (Xiao et al., 2006). This parameter quantifies the % of HL-60 cells with at least one bound platelet.

### Animal studies

All animal studies were approved by the Roswell Park Cancer Institute Animal Care and Use Committee (RPCI-IACUC). Mouse bone marrow cells (BMCs) were isolated from the tibia and femur of C57BL/6 mice. A histopaque gradient was used to obtain mouse polymorphonuclear leukocytes (PMNs) from these BMCs (Wang et al., 2018). Both the isolated BMCs and PMNs were independently cultured with either 50-80μM of Ac_5_GalNTGc/Ac_5_GlcNTGc/Ac_4_GalNAc or vehicle control (0.125-0.2% DMSO) in endotoxin-free IMDM with G-CSF, IL-3 and 10%FBS. At ~40h, the granulocytes were gated in the cytometer using nuclear stain LDS-751 and anti-mouse Ly-6G clone 1A8. Cell surface PSGL-1 expression (clone 2PH1), selectin-IgG binding (P-, E- and L-selectin) and VVA binding were quantified.

In some cases, Ac_5_GalNTGc and vehicle control treated BMCs were differentially tagged with cell tracker dyes CMFDA (5-chloromethylfluorescein diacetate, green) or CMTMR (5-(and-6)-(((4-chloromethyl)benzoyl)amino)tetramethylrhodamine, red) following manufacturer’s instructions (Setareh Biotech, Eugene, OR). Labeled cells were mixed at 1:1 ratio, and injected *i.v*. into recipient mice 1h after induction of peritonitis using 4% thioglycollate (Wang et al., 2018). 20h thereafter, peritoneum lavage and bone marrow cells were collected and analyzed using a BD LSR-II flow cytometer. APC conjugated anti-mouse Ly-6G 1A8 was used to identify granulocyte, and the ratio of red/green cells in the samples was quantified.

To test pharmacological efficacy, mice were injected with 100mg Ac_5_GalNTGc/kg/day or vehicle control for 4 days. Thioglycollate induced peritonitis was then induced as above for 16h. Granulocyte counts in the peritoneal lavage and bone marrow were quantified using anti-mouse CD11b, macrophage marker F4/80 and anti-Ly-6G (1A8) in cytometry runs. VVA binding to cells was also measured. Complete blood count (CBC) analysis was also performed using blood samples at the end point.

### Cellular O-glycome analysis

0.5×10^6^ HL60 cells were cultured in serum-free, phenol red-free Advance Dulbecco’s Modified Eagle Medium (ADMEM) along with 80μM Ac_5_GalNTGc or vehicle control for 16h. 100μM peracetylated GalNAc-O-Bn available from a previous study was then added to the cells (Wang et al., 2018). After 48h, GalNAc-O-Bn related products were purified from cell culture medium using Sep-Pak C18 cartridges (Waters, Milford, MA), permethylated and analyzed using ESI (electrospray ionization) LC-MS/MS (Orbitrap-XL mass spectrometer, Thermo). Instrument MS^1^ tolerance was 15ppm and MS/MS data were obtained following collision induced dissociation (30eV) at 1Da resolution using the ion-trap detector. All spectra are annotated using DrawGlycan-SNFG (Cheng et al., 2017).

### MALDI-TOF MS glycomics profiling

*N*-linked and GSL derived glycans were extracted from Ac_5_GalNTGc or vehicle control HL60s as described previously (Mondal et al., 2015). All glycans were permethylated prior to MALDI-TOF MS and MALDI-TOF-TOF MS/MS analysis. Released glycans from GSLs were deuteroreduced prior to permethylation. Data were annotated using the glycobioinformatics tool, GlycoWorkBench (Ceroni et al., 2008). The proposed assignments for the selected peaks were based on ^12^C isotopic composition together with knowledge of the biosynthetic pathways. The proposed structures were confirmed using MS/MS.

### β1,3GalT/C1GalT1 activity

5μL of HL-60 cell lysate (10mg/mL) were added to 15μL of a reaction mixture (500μM GalNAc-OBn, 1 mM UDP-Gal, 7 mM ATP and 20 mM Mn(OAc)_2_ in HEPES) and then incubated at 37°C overnight. After incubation, 1mL of 70% ACN was added to precipitate proteins, mixture vortexed and centrifuge-pellet removed (14,000g x 5 min). Supernatant collected was solvent evaporated and resuspended in 50μL of 50% MeOH (containing 10μM of 2OMe-Galβ1-3GlcNAc-OBn as internal standard; *m/z* [M+H]^+^ = 488.21264). For LC-MS/MS, 10μL sample was injected into Q Exactive™ Hybrid Quadrupole-Orbitrap Mass Spectrometer (Thermo) equipped with a C18 column (Waters Xselect CSH C18, 3.5μm, 2.1×150 mm). The mobile phases were A: water and B: acetonitrile (CH_3_CN), containing 0.1 % formic acid in both phases. Data were acquired over 95 min at a flow rate of 0.1 mL/min using the following gradient: (i) from 0% to 13% B (0-5 min); (ii) 13% to 30% B (5-85 min); (iii) 30% to 100% B (85-90 min); and (iv) 100% to 0% B (90-95 min). MS1 data were acquired using the Orbitrap detector (60,000 resolution) and MS/MS in HCD mode (30% collision energy). Data analysis was carried out by calculating the area under the curve of the XIC curve in triplicates, using XCalibur software (Thermo).

### Sugar nucleotide analysis

Nucleotide-sugar analysis was performed as described recently (Del Solar et al., 2020). Briefly, 1.2×10^6^ HL60 cells treated with 80μM Ac_5_GalNTGc, Ac_5_GlcNTGc, Ac_4_GalNAc or vehicle (0.2% DMSO) were resuspended in 75% ACN (containing 10 μM of CMP-Neu5Gc as internal standard; *m/z* [M-H]--= 629.13491) and spun down at 16,000g at 4°C for 10 min. Supernatant was collected and injected into LC-MS system. For LC-MS/MS analysis, 5 μL of sample were injected into the Q Exactive instrument described above, equipped with a HILIC column (Waters XBridge Amide 3.5μm, 2.1×150 mm). The mobile phases were, A: water containing 5mM NH_4_OH and 5mM NH_4_OAc and B: 90% acetonitrile (CH_3_CN), containing 5mM NH_4_OAc. Data were acquired over 60 min at a flow rate of 0.1 mL/min and 40 °C using the following gradient: (i) from 90% to 85% B (0-5 min); (ii) 85% to 80% B (5-45 min); (iii) 80% to 60% B (45-55 min); and (iv) 60% to 90% B (55-60 min). MS^1^ data were acquired using the Orbitrap detector (60,000 resolution) and MS/MS in HCD mode (28% collision energy). Analysis was carried out in triplicates.

## QUANTIFICATION AND STATISTICAL ANALYSIS

All error bars represent standard deviations for >3 repeats. Discrete data points in individual panels of main manuscript specify sample size. Student’s two-tailed t-test was performed for dual comparisons. Analysis of variance (ANOVA) followed by the Student-Newman-Keuls post-test was used for multiple comparisons. *P*<0.05 was considered to be statistically significant.

## DATA AND CODE AVAILABILITY

The data and reagents that support the findings of this study are available from the corresponding authors. Glycan sketches in this manuscript were generated using DrawGlycan-SNFG (https://github.com/neel-lab/DrawGlycan-SNFGv2; https:\\VirtualGlycome.org/drawglycan). MALDI-TOF data analysis was performed using GlycoWorkBench (https://github.com/alternativeTime/glycoworkbench). Both are open-source programs.

## Notes

### Competing Interest Statement

The authors have declared no competing interest.

### Summary of Updates

Spelling of one of the co-authors corrected. Rest of document is the same.

