## Supplemental Figures for "Efficient Inhibition of O-glycan biosynthesis using the hexosamine analog Ac_5_GalNTGc"

### Supplemental Material

**Contents**

Supplemental Figures S1-S7
Supplemental Tables S1-S2

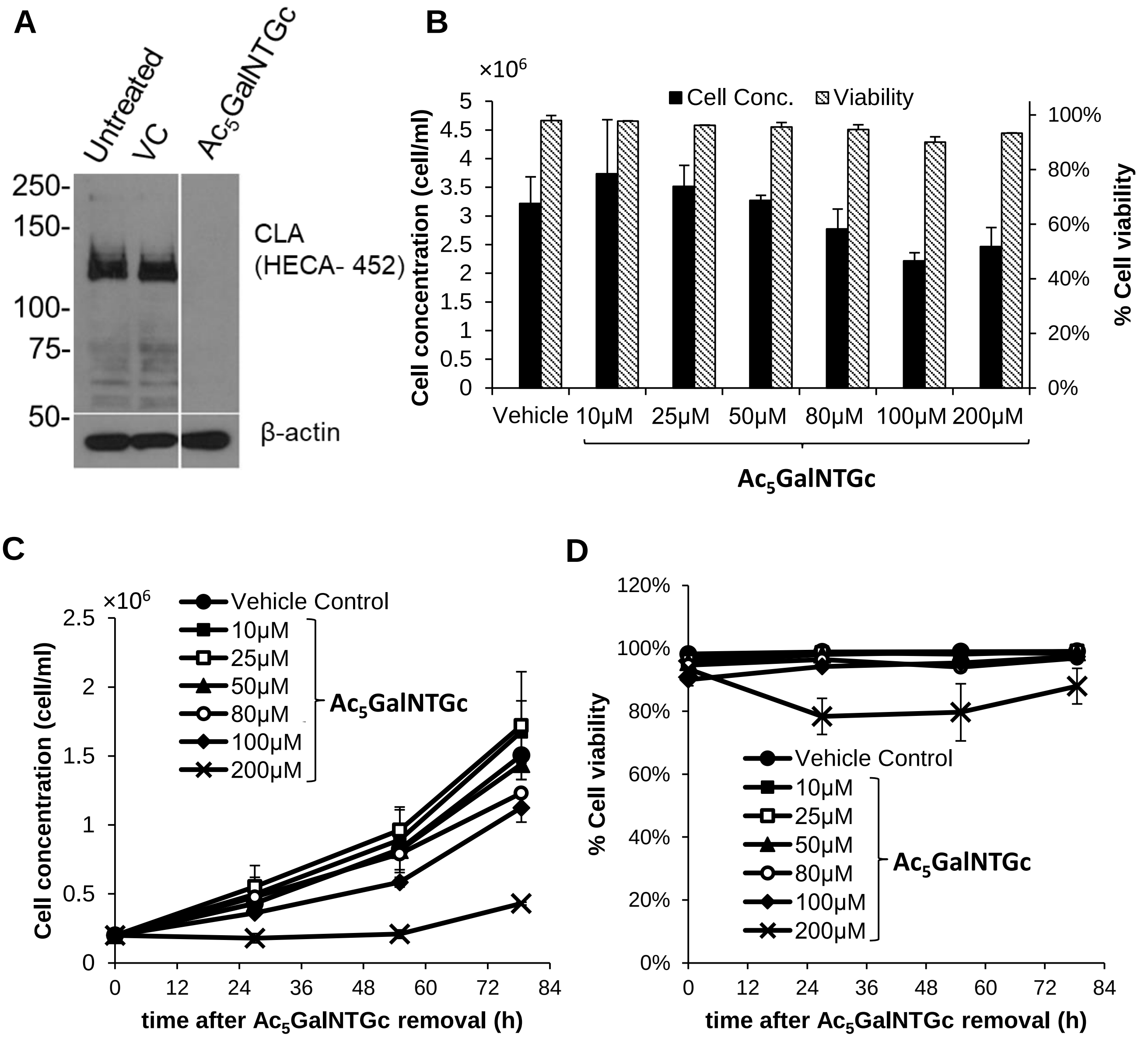


**Figure S1 (Related to Fig. 2). Effect of $Ac_5$GalNTGc on HL60 cell CLA (HECA-452) expression, cell viability and growth rate. A.** Western blot for CLA (HECA-452) in total lysates of HL-60 cells incubated with $Ac_5$GalNTGc (100 μM, 48 h) along with untreated and vehicle-treated controls. β-actin blots were used as loading controls. Blots shown are representative of at least two biological replicates. **B.** $0.5\times10^6$ HL60 cells/mL were cultured with 0-200μM $Ac_5$GalNTGc (Vehicle control= 0.5% DMSO, corresponding to 200μM $Ac_5$GalNTGc). Cell concentration was measured using a hemocytometer and viability based on trypan-blue exclusion. **C-D.** Following culture of HL60s for 40 h with various concentrations of $Ac_5$GalNTGc, the cells were washed to remove the small molecule. The washed cells were then re-suspended in fresh growth medium at $0.2\times10^6$ cells/mL for additional 3-4 days. Cell concentration (C) and viability (D) were continuously monitored. The cell growth rate and viability were not altered by $Ac_5$GalNTGc below 100μM. Cell growth lag phase was higher at 200μM.

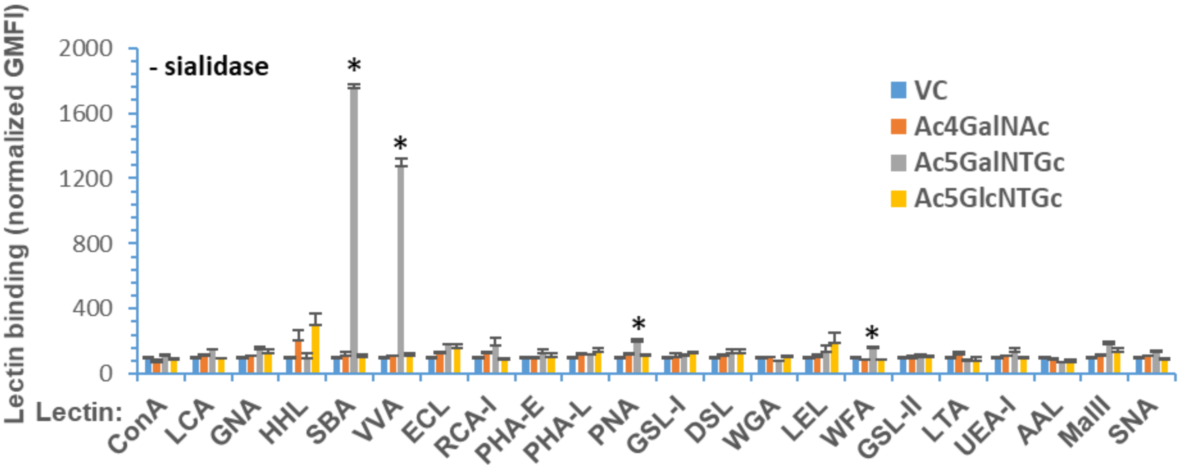


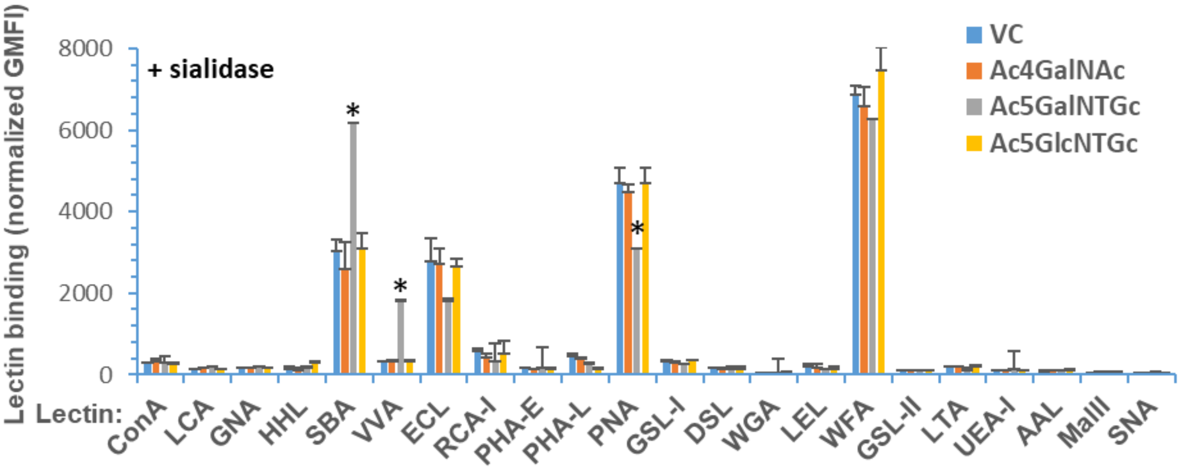


| Lectin | Binding specificity | Lectin | Binding specificity |
|---|---|---|---|
| ConA | a(Man) in high mannose | PHA-L | Gal(β1-4)GlcNAc(β1-2/6)Man compelx |
| LCA | a(Man) with α(1-6)Fuc dependence | PHA-E | Gal(β1-4)GlcNAc(β1-2)Man with bisect |
| GNA | Man(a3)Man | MAL-II | Neu5Ac(α2-3)Gal |
| HHL | α(1-3) & α(1-6) linked Man | SNA | Neu5Ac(α2-6) linked glycans |
| SBA | GalNAc(α/β) | AAL | Fuc(α1-3/6) and also Fuc(α1-2) |
| VVA | GalNAc(α/β) | UEA-I | Fuc(α1-2) [blood gp.] |
| PNA | Galα(1-3)GalNAc in O-glycans | LTA | Fuc(α1-2) |
| ECL | Gal, lactose based glycans | GSL-I | GalNAcβ on B-gp |
| RCA-I | Gal, lactose | GSL-II | GlcNAcβ |
| WFA | Gal(β1,4)GlcNAc, but also GalNAc(α/β1,3/6)Gal | LEL | GlcNAc terminal on endothelial cells, also Gal terminated sturctures |
| DSL | GlcNAcβ(1-4/6) linked | WGA | GlcNAc, oligomers, and also chitobiose core |

**Figure S2 (Related to Fig. 3). Lectin staining following $Ac_5$GalNTGc treatment.** HL60 cells were cultured with either vehicle control or one of the peracetylated HexNAc analogs (80µM, 40h). Cells were then probed with a panel of fluorescent lectins, either prior to (top) or following (bottom) sialidase treatment. Data are Mean $\pm$ SD. MFI normalization was performed by setting MFI of vehicle control for each lectin, without sialidase, to 100. This simplifies comparison of treatment effects. Table lists lectin binding specificity based on Consortium of Functional Glycomics microarray data and literature sources. $Ac_5$GalNTGc only affected the binding of lectins recognizing O-glycan epitopes (SBA, VVA, PNA).

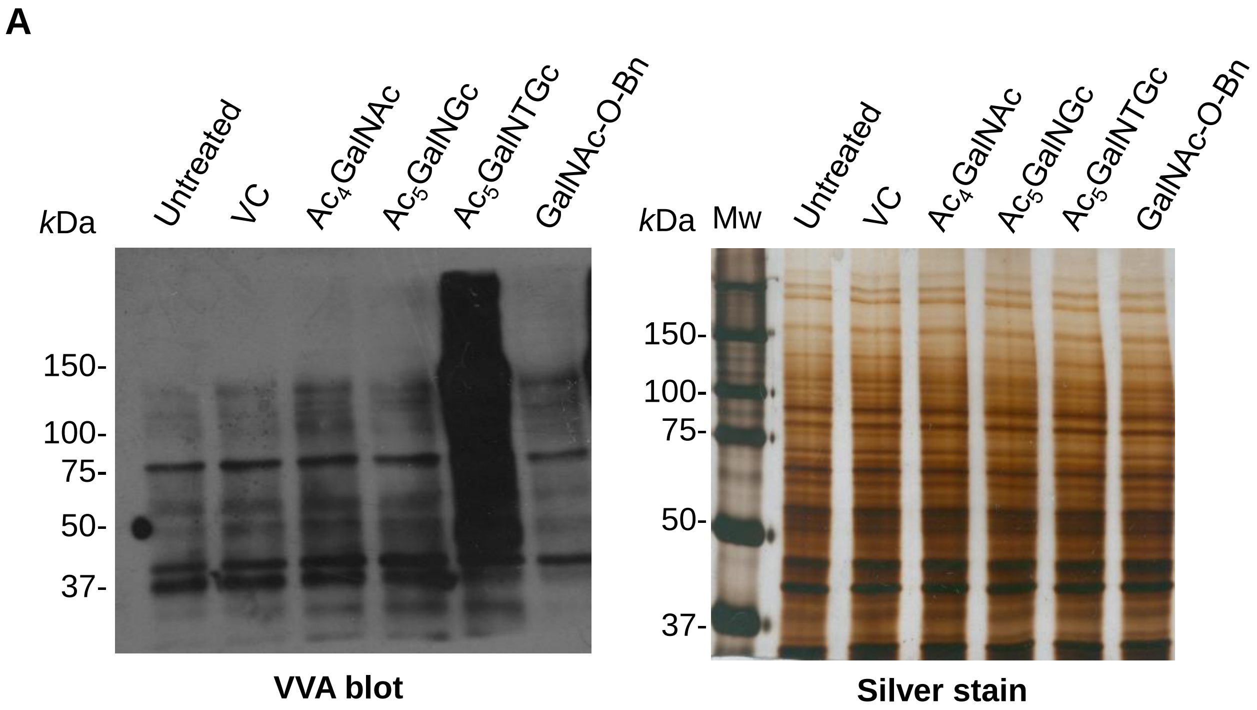


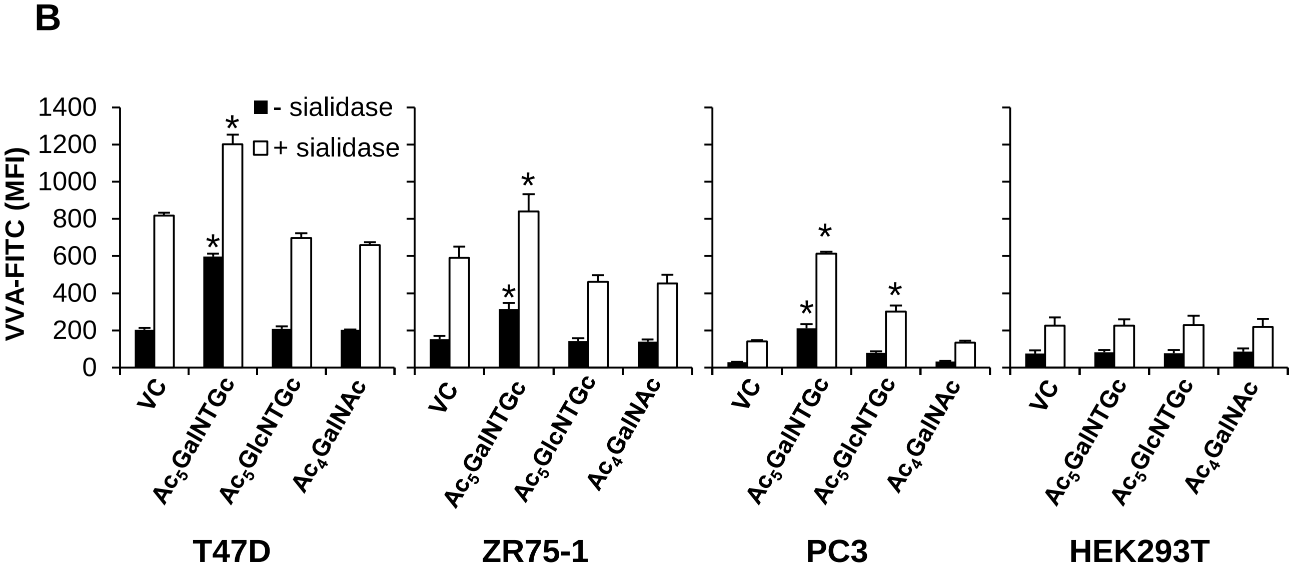


**Figure S3 (Related to Fig. 3). $Ac_5GalNTGc$ augments VVA-lectin binding in heterologous cell types. A**. *Vicia villosa* agglutinin (VVA) blot of total cell lysates of HL-60 cells incubated with peracetylated GalNAc analog (100 mM, 48 h). Biotinylated-VVA was employed followed by HRP-avidin staining. Silver stained gels are shown as loading controls. Blots and gels are representative of at least two biological replicates. **B**. 80µM $Ac_5GalNTGc$, $Ac_5GlcNTGc$ and $Ac_4GalNAc$ were added to the culture medium of T47D (breast), ZR75-1 (breast), PC3 (prostate), and HEK293T (kidney) cells at $0.5\times10^6$ cells/ml. At 40h, VVA-lectin binding, measured by flow cytometry, was augmented in all cell types except HEK293Ts suggesting that $Ac_5GalNTGc$ may truncate O-glycan biosynthesis in many different cell types. * $P$<0.05 with respect to other treatments under the same category (i.e. sialidase to sialidase, no sialidase to no sialidase).

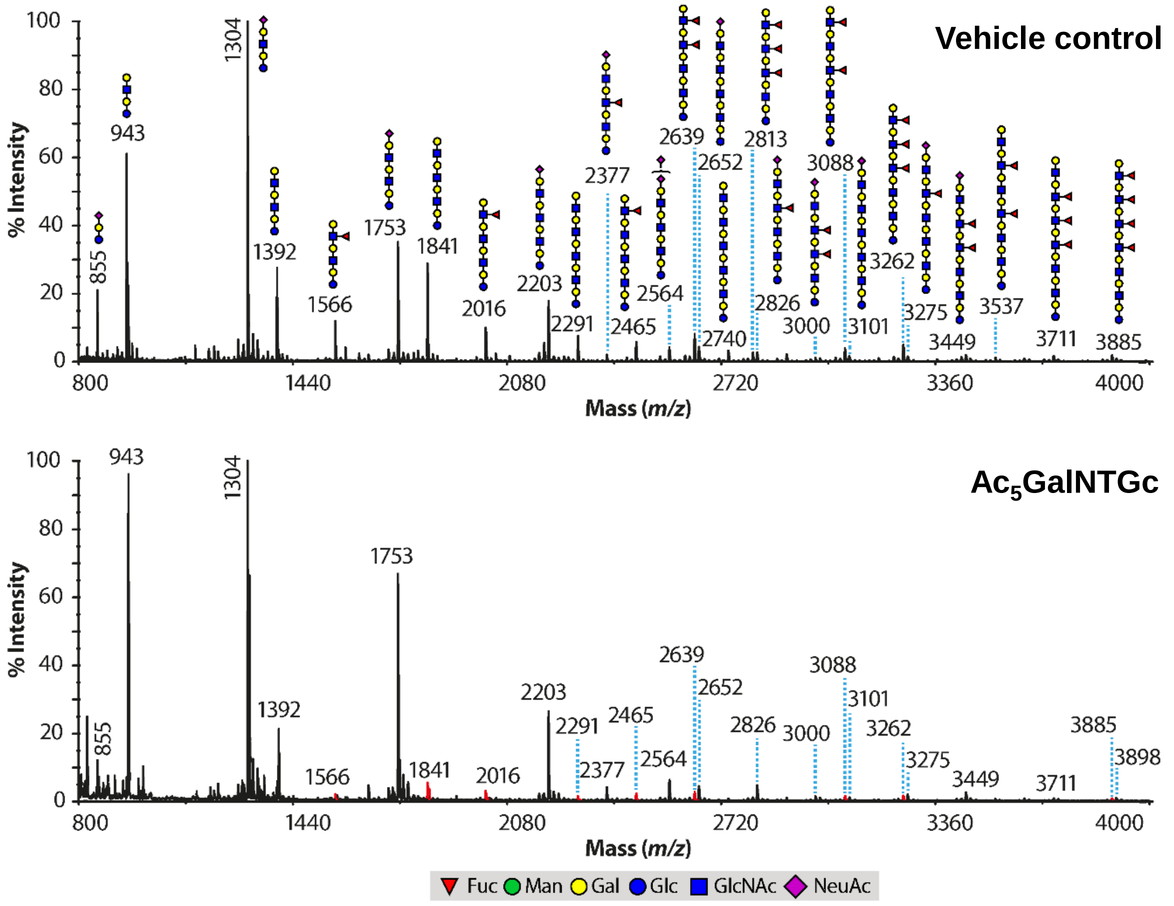


**Figure S4 (Related to Fig. 6). Effect of $Ac_5$GalNTGc on GSL derived glycans.** MALDI-TOF MS profile of permethylated glycans derived from GSLs for cells treated with 80μM $Ac_5$GalNTGc (lower panel) or vehicle (upper panel) for 40h. Putative structures are based on composition, tandem MS, and knowledge of biosynthetic pathways. All molecular ions are $[M+Na]^+$.

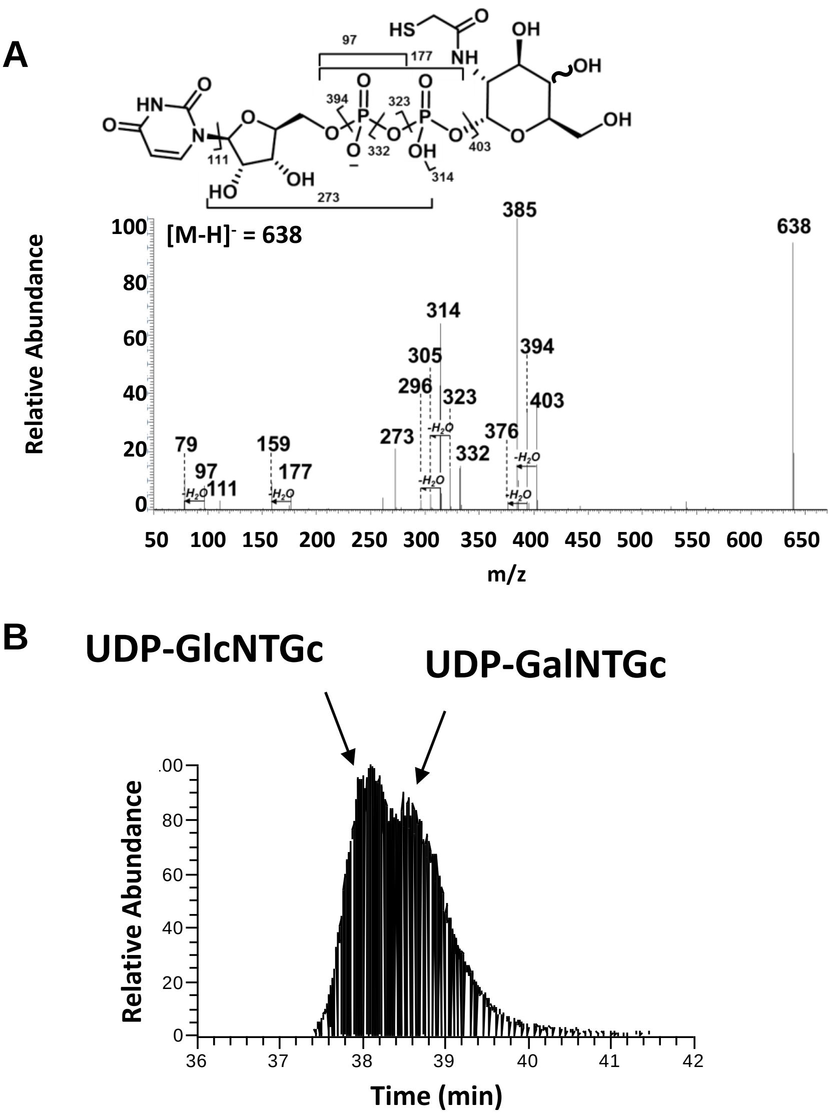


**Figure S5 (Related to Fig. 7). Nucleotide-sugar analysis: A.** MS/MS fragmentation profile of UDP-HexNTGc. Distinction between UDP-GlcNTGc and UDP-GalNTGc is not possible due to absence of cross-ring fragmentation. **B.** UDP-HexNTGc elutes as two overlapping peaks, which presumably include UDP-GlcNTGc and UDP-GalNTGc as indicated. Similar overlapping elution profiles, with Glc appearing just prior to Gal, were observed both for mixtures of synthetic UDP-Glc/UDP-Gal, and UDP-GlcNAc/UDP-GalNAc standards.

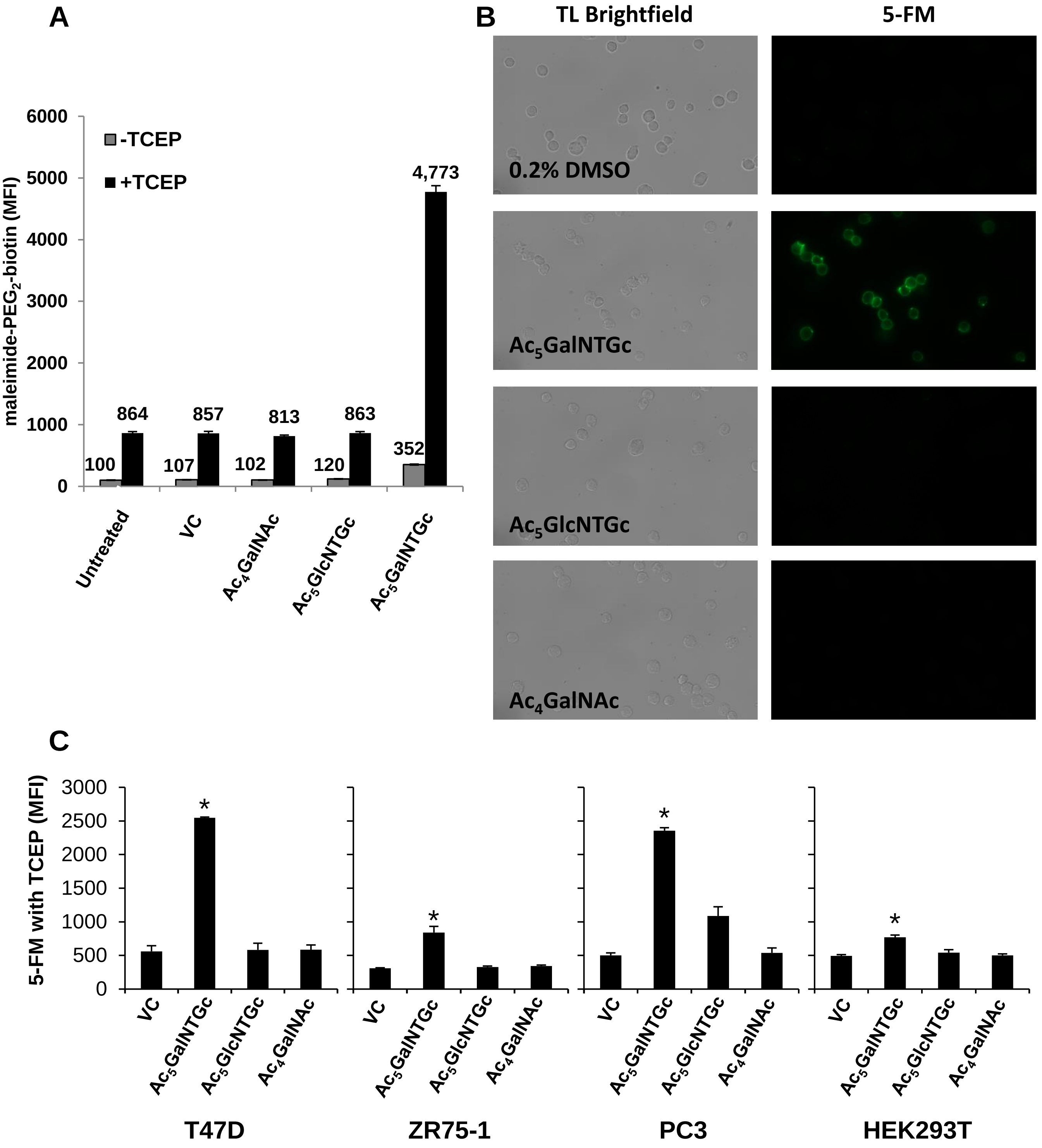


**Figure S6 (Related to Fig. 7) 5-FM binding on HexNAc treated cells. A.** HL-60 cells cultured with peracetylated HexNAc analogs (100 μM, 48 h) were labeled using Michael addition reaction of maleimide-PEG$_2$-biotin to accessible free suldfhydryl groups on the cell surface before (-TCEP) and after (+TCEP) treatment, followed by FITC-conjugate avidin staining. **B.** 0.5×10$^6$ HL60 cells/mL were cultured with VC (0.2% DMSO), 80μM of Ac$_5$GalNTGc, Ac$_5$GlcNTGc or Ac$_4$GalNAc for 40h. 5-FM only reacted with cells treated with Ac$_5$GalNTGc. **C.** 80μM Ac$_5$GalNTGc, Ac$_5$GlcNTGc and Ac$_4$GalNAc were added to the culture medium of T47D, ZR75-1, PC3, and HEK293T. 5-FM coupling to cells at 40h was assessed using flow cytometry. 5-FM incorporation levels varied with cell type, and was lowest for HEK293T cells. * $P$<0.05 with respect to other treatments for the same cell type.

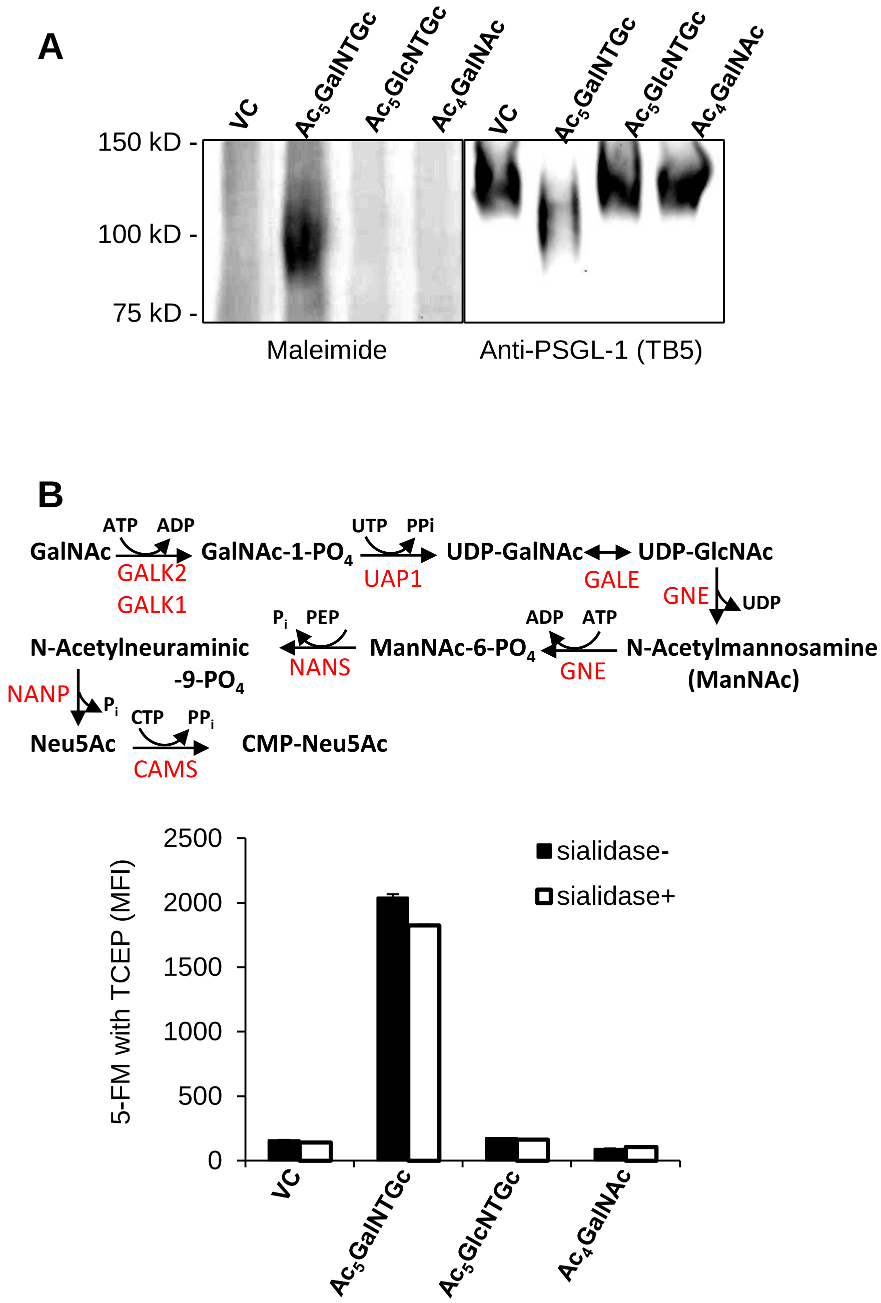


**Figure S7 (Related to Fig. 7) Direct incorporation of GalNTGc.** HL60s cultured with 80μM $Ac_5$GalNTGc, $Ac_5$GlcNTGc, $Ac_4$GalNAc or VC were labeled with maleimide-$PEG_2$-biotin (**A**) or maleimide-FITC (**B**). **A.** PSGL-1 was immunoprecipitated from equal amounts of cells and resolved using 4-20% SDS-PAGE under reducing conditions. Detection was performed using anti-biotin for thiol (left panel) or anti-PSGL-1 clone TB5 (right panel). $Ac_5$GalNTGc treated samples reacted well with anti-biotin suggesting maleimide incorporation into PSGL-1. PSGL-1 molecular mass was also reduced by $Ac_5$GalNTGc. **B.** Reaction pathway shows the biochemical steps and enzymes (red) required for the conversion of GalNAc (or GalNTGc) to CMP-sialic acid (or modified sugar-nucleotides). HL60 cells cultured with modified monosaccharides were labeled with 5-FM in the presence of TCEP. A portion of the cells were then treated with α2-3,6,8,9 *Arthrobacter Ureafaciens* neuraminidase to remove sialic acid. Maleimide incorporation measurements made before and after sialidase treatment, suggest that $Ac_5$GalNTGc may not be incorporated nor convert into sialic acids. Similar results were observed even when cells were sialidase treated prior to 5-FM labeling. It is assumed here that the neuraminidase can cleave sialic acid with thiol substituents.

**Table S1. Complete blood count for mouse treated with inhibitor**

| | $Ac_5GalNTGc$ | | Control | | |
|---|---|---|---|---|---|
| **Samples** | **Average** | **±SEM** | **Average** | **±SEM** | **Significance** |
| **WBC (cell/μl)** | 4,897.5 | 1,188.09 | 4,555 | 959.88 | n.s. |
| **PMN** | 13.53% | 1.51% | 10.85% | 1.02% | n.s. |
| **Lymphocyte** | 76.88% | 2.56% | 81.43% | 1.70% | n.s. |
| **Monocyte** | 2.50% | 0.14% | 2.53% | 0.33% | n.s. |
| **Eosinophil** | 4.23% | 0.64% | 1.93% | 0.58% | n.s. |
| **Basophil** | 0.98% | 0.38% | 0.83% | 0.20% | n.s. |
| **RBC (cell/μl)** | 9,452,500 | 30,923.29 | 9,517,500 | 228,669.2 | n.s. |
| **PLT (cell/μl)** | 1,056,750 | 51,002.25 | 1,000,750 | 116,401.2 | n.s. |

**(Related to Fig. 5)** CBC was performed 4-days after systemic $Ac_5$GalNTGc infusion. $Ac_5$GalNTGc did no affect platelet, red blood cell, total leukocyte or leukocyte differentials. n.s. : not significant.

**Table S2: Gene expression data provided as transcript per million from CCLE database***

| | HEK-kidney | HL60-leukocyte | PC3-prostate | T47D-breast | ZR751-breast |
|---|---|---|---|---|---|
| **core-1 synthase** | | | | | |
| C1GALT1 | 19.73 | 3.31 | 21.58 | 8.95 | 20.97 |
| C1GALT1C1 | 37.19 | 10.86 | 34.88 | 22.73 | 41.79 |
| **ppGalNAcTs** | | | | | |
| GALNT1 | 38.77 | 27.3 | 21.61 | 19.93 | 72.52 |
| GALNT2 | 97.14 | 22.88 | 104.81 | 30.45 | 22.34 |
| GALNT3 | 28.35 | 7.79 | 34.74 | 11.16 | 55.63 |
| GALNT4 | 2.19 | 0.56 | 4.76 | 4.12 | 3.72 |
| GALNT5 | 1.37 | 0.06 | 0.01 | 0.2 | 0.15 |
| GALNT6 | 11.77 | 5.44 | 25.91 | 55.99 | 57.81 |
| GALNT7 | 44.25 | 9.52 | 8.71 | 11.97 | 34.57 |
| GALNT8 | 0.01 | 0.05 | 0 | 0 | 0 |
| GALNT9 | 0.1 | 0 | 0.26 | 0.09 | 0 |
| GALNT10 | 57.88 | 2.57 | 27.99 | 50.36 | 32.16 |
| GALNT11 | 16.84 | 9.61 | 15.17 | 22.2 | 35.65 |
| GALNT12 | 3.87 | 0.11 | 6.13 | 2.04 | 4.74 |
| GALNT13 | 0.14 | 0 | 0.47 | 0.01 | 0.13 |
| GALNT14 | 38 | 29.6 | 2.69 | 0.38 | 5.5 |
| GALNT15 | 0.29 | 0 | 0.02 | 0 | 0.01 |
| GALNT16 | 0.7 | 0.05 | 0.28 | 0.26 | 1.82 |
| GALNT17 | 0 | 0 | 0 | 0.01 | 0 |
| GALNT18 | 1.14 | 0.34 | 15.46 | 8.49 | 8.29 |
| **Enzymes involved in salvage pathway** | | | | | |
| GALK1 | 21.78 | 34.2 | 23.98 | 34.74 | 44.9 |
| GALK2 | 13.74 | 10.88 | 11.96 | 30.65 | 12.02 |
| UAP1 | 104.34 | 26.26 | 83.14 | 24.54 | 51.75 |
| GALE | 67.52 | 21.99 | 49.99 | 63.51 | 85.92 |
| **selected sialyltransferases** | | | | | |
| ST3GAL1 | 10.89 | 5.94 | 50.51 | 11.7 | 20.23 |
| ST3GAL4 | 17.14 | 58.08 | 54.33 | 53.29 | 43.17 |
| ST6GALNAC1 | 0.02 | 0.01 | 1.13 | 0.21 | 0.32 |
| ST6GALNAC2 | 0.58 | 0.48 | 1.7 | 50.89 | 86.87 |
| ST6GALNAC4 | 10.42 | 2.63 | 6.27 | 5.18 | 51.53 |

* CCLE: https://portals.broadinstitute.org/ccle; data include sum of all transcript variants
